# Hepatic ADMA–PRMT1 axis regulation is associated with NO-dependent endothelial dysfunction in MASH

**DOI:** 10.1101/2025.06.10.658894

**Authors:** Justine Lallement, Sebastian Bott, Hrag Esfahani, Kristian Serafimov, Maxence Henderson, Zoé Benoit, Laurent Dumas, Sultan Al-Siyabi, Olivier Feron, Isabelle Anne Leclercq, Chantal Dessy

## Abstract

Metabolic dysfunction-associated steatohepatitis (MASH) is a severe form of fatty liver disease and a recognized cardiovascular risk factor, yet the mechanisms linking hepatic pathology to vascular dysfunction remain poorly understood. We aimed to investigate whether MASH impairs endothelial function via nitric oxide (NO)-dependent mechanisms and to identify potential liver-derived mediators involved in this process.

Endothelial function was assessed in two murine MASH models, Foz mice fed a high-fat diet and C57BL/6JRj mice fed a western diet with fructose, by using wire myography, while blood pressure was monitored via telemetry. NO pathway was further investigated through eNOS expression and activation and Hb-NO measurements. ADMA metabolism was analyzed in both liver tissue and plasma by LC-MS and gene expressions. Additionally, bovine aortic endothelial cells (BAECs) were treated with mouse plasma to measure the circulating factors’ effects on eNOS activation and the role of oxidative stress.

Both models exhibited impaired NO-dependent vasorelaxation without evidence of atherosclerosis. In Foz mice, this impairment was associated with reduced eNOS expression and activation. Surprisingly, plasma Hb-NO levels did not reflect vascular NO deficiency, likely due to elevated hepatic *iNOS* expression. In both models, hepatic and plasma ADMA levels were increased, concomitant with hepatic upregulation of *Prmt1*. BAECs exposed to plasma from MASH mice showed reduced eNOS activation independent of oxidative stress.

Our findings reveal that MASH is consistently associated with NO-dependent endothelial dysfunction, with ADMA emerging as a key liver-derived mediator. The PRMT1/ADMA/NO axis may represent a mechanistic link between liver pathology and vascular impairment, positioning ADMA as a potential biomarker and therapeutic target for cardiovascular risk associated with MASH.

**Highlights:**

- MASH is associated with impaired NO-dependent endothelial function in two distinct MASH models.
- Circulating factors disrupt the NOS/NO pathway independently of oxidative stress
- Reduced plasma Hb-NO levels do not accurately reflect NO-dependent endothelial dysfunction in a context of MASH.
- Elevated plasma and hepatic ADMA levels associated with hepatic *prmt1* upregulation are consistently observed in both MASH models.
- The PRMT1/ADMA/NO axis emerges as a key liver-mediated mechanism driving endothelial dysfunction in MASH

## Introduction

Metabolic dysfunction-associated steatotic liver disease (MASLD) currently affects about 30% of the adult population worldwide^1^. Recently, MASLD has been redefined as steatotic liver associated with at least one of the following cardiometabolic risk factors: (1) increase in body mass index or waist circumference; (2) impaired glucose metabolism; (3) high blood pressure; (4) high triglyceride levels; (5) low high-density cholesterol levels^2^. Between 10 and 30% of patients will progress toward metabolic dysfunction-associated steatohepatitis (MASH), the aggressive from of the disease, with a potential to evolve towards severe fibrosis, cirrhosis and hepatocellular carcinoma^3^. Besides liver-related morbidity and mortality, MASLD and MASH patients are at higher risk of developing cardiovascular disease (CVD) than those without. Notably CVD stands as the leading cause of morbi- mortality in patients with MASLD^4–6^. While MASLD and CVD share several well-established risk factors, such as obesity, insulin resistance, and low-grade inflammation, it remains challenging to determine whether MASLD acts as a primary driver or an effective an effective contributor of CVD. Even more, the literature reveals that the progression of liver disease may exacerbate CVD, and vice versa^7^. However, large cohort studies increasingly support the view that MASH is an independent risk factor for CVD, rather than simply a bystander or outcome of shared metabolic conditions^5,8,9^.

The primary cause of MASLD is an imbalance in lipid metabolism, driven by both dietary lipid overload and insulin resistance (IR)^10^. In adipose tissue, IR promotes uncontrolled lipolysis in adipocytes, leading to an excessive influx of free fatty acids (FFAs) into the liver^11^. Concurrently, hepatic *de novo* lipogenesis is enhanced under these conditions, further increasing the FFA burden. FFA accumulation drives excessive β-oxidation in mitochondria, resulting in increased ROS production and consequently oxidative stress^12^. This lipid overload and oxidative imbalance contribute to hepatocellular injury, inflammation, and ultimately, MASLD progression^13^. Notably, these same mechanisms, oxidative stress, dyslipidemia and inflammation, are also considered as key drivers of endothelial dysfunction, an early mechanism in the development of atherosclerosis and hypertension leading to CV complications^14–16^.

MASLD/MASH patients exhibit significantly reduced flow-mediated vascular dilation compared to healthy or obese controls, suggesting that hepatic metabolic disturbances drive endothelial dysfunction^17^. Importantly, patients with MASH display a markedly diminished vascular response relative to individuals with simple steatosis^18,19^ primarily attributed to alterations in the nitric oxide (NO) pathway^20^. Endothelial NO, synthesized by endothelial nitric oxide synthase (eNOS), critically regulates vascular tone and inhibits leukocyte adhesion and platelet aggregation, thereby maintaining overall vascular homeostasis. Perturbations in NO bioavailability are a hallmark of early endothelial dysfunction and a predictor of atherosclerotic disease^21,22^. NO bioavailability is tightly regulated through multiple mechanisms, including the modulation of eNOS activity and the availability of its substrate, L-arginine. Under oxidative stress and substrate depletion, conditions encountered in MASLD/MASH patients, eNOS could become "uncoupled" and generate superoxide instead of NO. This paradoxical shift not only would reduce NO bioavailability but also contribute directly to vascular oxidative damage^23^. Furthermore, increased levels of asymmetric dimethylarginine (ADMA), an endogenous inhibitor of eNOS, competitively inhibits L-arginine binding to eNOS, further reducing NO production. Moreover, both hypercholesterolemia and inflammation were shown to increase ADMA accumulation, particularly in the liver^24,25^. Together, these findings suggest that dyslipidemia, systemic oxidative stress, and inflammation associated with liver disease could act synergistically to drive endothelial dysfunction in MASH, thereby contributing to an increased risk of cardiovascular disease.

In this study, we examine vascular function in two well-established MASH mouse models in which we previously demonstrated pathological cardiac remodeling linked to MASH disease^26^, we aim to explore endothelial function in association with MASH, and investigate the molecular mechanisms at play as to unravel liver-driven contributors to endothelial dysfunction.

## Materials and methods

### Animals and diets

The fat Aussie mouse strain (Foz), which carries a truncating mutation in the Alms1 gene on a non-obese diabetic (NOD.B10) background, was bred in our animal facility under controlled 12-hour light/dark cycles with unlimited access to water and food. Male homozygous mutants and their homozygous wild-type littermates (referred to as WT) were turned on a high-fat diet (HFD, catalog n°D12492 - 60 kcal% from fat, Research Diets, NJ, USA) at 8 weeks of age or maintained on standard rodent chow (normal diet [ND], SAFE® A03 - 13.5 kcal% from fat, SAFE SAS, France) for 24 weeks.

C57BL/6JRj male mice, purchased from Janvier-Labs (France) at 7 weeks of age, were randomized on different diets at 8 weeks: high-fat diet (HFD, catalog n° D12492 - 60 kcal% from fat, Research Diets, NJ, USA), high-fat diet supplemented with 30% fructose in drinking water, western diet (WD, catalog n° D12079B - 40 kcal% from fat, 40% kcal from sucrose, 0.5% cholesterol, Research Diets, NJ, USA) supplemented with fructose in drinking water, for 24 weeks. One group of mice was also maintained on rodent chow as healthy controls (normal diet [ND] - SAFE® A03 - 13.5 kcal% from fat, SAFE SAS, France). Animal experimentation adhered to regulatory guidelines for the humane care of laboratory animals as established at UCLouvain, in accordance with European regulations. The study protocol was approved by UCLouvain’s ethics committee (2016/UCL/MD/003, 2020/UCL/MD/019, 2021/MD/02 and 2022/UCL/MD/62).

### Histology for NAFLD activity score

Routine histological evaluation for NAFLD activity score (NAS) assessment were performed on 5 µm thick sections of paraffin-embedded livers stained with haematoxylin & eosin (H&E), following the method described by Kleiner et al^27^. This approach, previously used to assess MASLD severity in this mouse strain^26^, considers a score above 5 as indicative of MASH. For immunohistochemical detection of F4/80, paraffin sections were treated with proteinase K, exposed to a primary rat anti-mouse F4/80 monoclonal antibody (1/200) (catalog n°MCA497G, BioRad, CA, USA). A rabbit anti-rat immunoglobulin (1/200) (catalog n°AI-4001, Vector laboratories, CA, USA) was then applied followed by a goat anti-rabbit streptavidin horseradish peroxidase-conjugated antibody (catalog n° K4003 EnVision, Agilent technologies, CA, USA). The peroxidase activity was revealed with diaminobenzidine (DAB, Agilent technologies, CA, USA) and slides were counterstained with haematoxylin. In addition to score the extent of hepatic inflammation, these specimens were also used for identification of crown-like structures, a morphological characteristic in progression from simple steatosis to MASH. Hepatic fibrosis was assessed on Sirius red (SR)-stained liver sections, using QuPath software (versions 0.3.0 & 0.4.3, Dr. Peter Bankhead, Edinburgh, UK) to determine the total tissue collagen content.

### Telemetry

Mice underwent surgery for implantation of a miniaturized telemetric device (PA-C10 Implant, Data Science International, MN, USA) after 22 weeks of diet. Using isoflurane-induced anesthesia a catheter was inserted and fixed in the left common carotid artery to enable remote real-time monitoring of blood pressure and heart rate in unrestrained animals. Prior surgery the mice received analgesic treatment (Temgesic® [buprenorphine], 0.1 mg/kg BW). After surgery, mice spent a week recovering. Blood pressure was recorded during three separate 24-hour periods (with a 24-hour interruption between measurements) both at baseline and after seven days of L-NAME treatment (Nω-Nitro-L-arginine methyl ester hydrochloride, catalog no. N5751, Sigma-Aldrich, Merck, Germany), administered in drinking water at 2 g/L. Data analysis was performed using Ponemah software (version v6.30, Data Science International, MN, USA). All surgeries, as well as data recording and analysis were performed by the same experienced operator (HE), blinded to the experimental groups.

### Wire myograph

The experiments were performed using a technique previously developed by the group^28,29^. At the time of sacrifice (terminal anesthesia), the mesenteric artery was harvested and cleaned from adherent tissues. Arterial rings (1.4–2 mm in length) from first-order mesenteric arteries (Foz model) or superior mesenteric arteries (C57BL/6JRj model) were mounted in myograph chambers (610M/620M; DMT, Denmark) filled with cold (4°C), low potassium chloride (5,9mM KCl) Krebs-Henseleit solution (hereafter designated as Krebs-buffer, see Table 2. supplementary methods) and oxygenated with a mix of O2 95% and CO2 5% (Air Liquide, Belgium). Chamber heating units were turned on to progressively (30min) accommodate the vessel segments to 37°C. During normalization, tension on each ring was adjusted to meet 90% of the vessel circumference developed by a fully relaxed vessel ring under a transmural pressure of 100mmHg, followed by 20min of resting. Vessel segments were then contracted for 10min with high-KCl (50mM) Krebs-solution. Those contracting with less than 1mN/mm were discarded from analysis. Properly reacting vessel rings were then washed three times with regular Krebs buffer and rested for 10min before measurements of vasodilative pathways. Each measurement comprised the induction of contraction in response to KCl-buffer (at 20 mM for C57BL/6Rj mice and 50 mM for Foz mice) and/or phenylephrine (Phe 10^-6^ or 10^-7^ M, catalog n°P1240000, Sigma-Aldrich, Merck, Germany) until stable contraction was reached, followed by examination of vasodilation capacity in response to carbachol (catalog n°PHR1511, Sigma-Aldrich, Merck, Germany) added in increasing concentrations (10^-7^ to 10^-5^M) with a 3min time interval between incremental doses^30^. We then used L-NAME (catalog n°N5751, Sigma-Aldrich, Merck, Germany), indomethacin (catalog n°405268, Sigma-Aldrich, Merck, Germany), or KCl high concentration to analyze separately relaxation mechanisms dependent on NO, prostacyclin (PGI2) and endothelium-derived hyperpolarizing factor (EDHFs), respectively. Next to prostaglandins with vasorelaxation properties, thromboxane A2 (TxA2), another arachidonic acid metabolite, has opposite effects. To unmask a possible TxA2-induced vessel contraction, we used the synthetic thromboxane receptor analog U46619 (dose: 1x10^-5^M, catalog n° BML-PG023-0001, Enzo Life Sciences, NY, USA). By saturating its target prior to measurement of PGI2-dependent relaxation, we prevented potential interference of TxA2 with our analysis. After each measurement, the chambers were rinsed three times with Krebs buffer and samples then rested for 10min before the next analysis. All measurements were monitored and recorded via LabChart 8 software (v8.1.16/AD Instruments, New Zealand). To maintain constant conditions during measurements, all buffers were preheated and bubbled with O2/CO2 to maintain the pH at 7.4 before application.

### Measurement of nitrosylated hemoglobin (HbNO) and asymmetric dimethylarginine (ADMA)

HbNO was measured by electron paramagnetic resonance (EPR) (5-coordinate-α-HbNO) as described^31^. ADMA in the plasma of Foz/WT mice was quantified using Fast ELISA (DLD Diagnostika, Germany) according to manufacturer’s specifications. Arginine, ADMA, citrulline and ornithine levels in the liver or plasma were measured by LC-MS after samples extraction and purification^32^.

### Liquid chromatography-mass spectrometry (LC-MS)

LC-MS analysis was performed on a Thermo Fisher Vanquish Horizon system equipped with a binary pump, autosampler and thermostated column compartment coupled to an Orbitrap Exploris 240 mass spectrometer^32^. Chromatographic separation was performed on a Waters (Antwerp, Belgium) Acquity Premier VanGuard-FIT BEH Amide column (100 x 2.1 mm, 1.7 µm), equipped with a VanGuard-FIT BEH Amide pre-column (5 x 2.1 mm, 1.7 µm)^32^. For metabolite analysis in ESI^+^ mode, a stock solution of 500 mM formic acid (catalog n°28905 Thermo Fisher Scientific, MA, USA) was prepared and adjusted to a pH of 3.5 with ammonium hydroxide (catalog n°105428, Sigma-Aldrich, Merck, Germany). A following 1:10 dilution with water (mobile phase A) and with acetonitrile (catalog n° 047138-K2, Thermo Fisher Scientific, MA, USA - mobile phase B) resulted in the final mobile phase conditions used during sample analysis in positive ionization mode. The chromatographic gradient was as follows: (0.0 min, 100% B; 20 min 70% B; 21 min 70% B; 21.01 min 100% B; 25 min 95% B) and a flow rate of 0.2 mL min^-1^ was used. A constant column temperature of 60 °C was maintained throughout the entire analytical run and the injection volume was 3 µL. The autosampler was kept at 4°C. Ion source parameters were as follows: Sheathe gas (35, nitrogen), aux gas (7, nitrogen), sweep gas (0 psi, nitrogen), ion transfer tube temperature (325°C), ion source voltage +3500 V, vaporizer temperature (275°C). Analytes were recorded via a PRM scan with a mass resolving power of 30000 and the RF lens setting was at 70 %. An isolation window of 2 *m/z* was used. Microscans were set to 1, AGC-Target value was set to 1e^5^ with a Maximum Injection Time of 54 ms. Collision Energy was normalized to 30/50/70%. Data processing comprising peak integration and regression was performed with Skyline^33^. Matrix matched calibration was performed via LC-MS standard addition with 1/x^2^-weighted linear regression. Calibration curves were constructed by plotting the peak area of the analyte (Buchem B.V., Netherlands) versus the area of the isotopic labeled internal standard (U^13^C, U^15^N Arginine, Sigma-Aldrich Merck, Germany).

### qPCR Analysis

After mechanical processing (Precellys Evolution, Bertin Technologies, France), total RNA from mouse aorta or liver was extracted using the Maxwell RSC microRNA Tissue kit (Promega, WI, USA). RNA concentration was ascertained by measuring optical density at 260 nm on a spectrophotometer NanoDrop (Thermo Fisher Scientific, MA, USA). 1 µg of total RNA from each sample was reverse transcribed with High Capacity cDNA Reverse Transcription Kit (Applied Biosystems, Lithuania). Real time quantitative PCR amplification reaction was carried out in a Rotor-Gene Q (Qiagen, Germany) using SYBR^TM^ Select Master Mix (Applied Biosystems, Lithuania) and primer pairs (Invitrogen, MA, USA) listed in Table 1, supplementary methods. To compare the fold change in target gene expression, relative quantification was performed using the 2^-ΔΔCt^ method, with ND or WT HF used as controls as appropriate. Data were normalized to the expression of a housekeeping gene, being *Ywhaz* for aorta samples and *Ppia* (cyclophiline-a) for liver samples, whose invariance of expression has been verified across groups.

### Western Blot Analysis

Samples of frozen mouse aorta or BAEC cells were homogenized in lysis buffer containing protease and phosphatase inhibitors (protease inhibitor cocktail, catalog n°P8340, Sigma-Aldrich, Merck, Germany; phosSTOP phosphatase inhibitor cocktail tablets, catalog n° 4906837001, Sigma-Aldrich, Merck, Germany). After high-speed shaking in Precellys (Precellys Evolution, Bertin Technologies, France) with stainless steel beads, the homogenate was centrifuged at 10 000 rpm for 10 min at 4 °C, and the supernatant was collected in a new tube. After quantitative analysis (PierceTM BCA protein assay kit, catalog n°A55864, Thermo Fisher Scientific, MA, USA), equal amounts of the protein samples (20 μg of BAEC cells or 40ug of aortic extracts) were resuspended in Laemmli sample buffer 1X and separated in an 8% sodium dodecylsulfate (SDS) polyacrylamide gel system (BIORAD, Hercules, CA, USA). After transfer and membrane blockage in milk or BSA 5%, the nitrocellulose membranes (catalog n°GE10600003, Amersham^TM^, Merck, Germany) were incubated with specific antibodies overnight at 4 °C and then with the secondary antibody conjugated with peroxidase for 1h at RT (Jackson Immunoresearch, MD, USA). The primary antibodies used were the following: anti-phospho-eNOS (Ser^1177^) (catalog n°9571S, 1/1000, Cell Signaling technology, MA, USA), anti-phospho-eNOS (Thr^495^) (clone 31 catalog n°612706, 1/1000, BD Biosciences, NJ, USA), anti-phospho-Akt (Thr^308^) (catalog n°9275, 1/1000, Cell Signaling technology, MA, USA), anti-eNOS (clone 3 catalog n°610297, 1/1000, BD Biosciences, NJ, USA), anti-Akt (catalog n°9272, 1/1000, Cell Signaling technology, MA, USA), anti HSP90 (catalog n°610419, 1/5000, BD Biosciences, NJ, USA). Immunoreactivity was detected in dark room or with Amersham^TM^ imager 680 (GE HealthCare, IL, USA) using an ECL detection reagent (catalog n°RPN2134, GE HealthCare, Il, USA or catalog n°WBKLSO500, Merck Milipore, MA, USA) and analyzed using the Image J software (LiCor Biosciences, NE, USA). Phosphorylated forms of Akt or eNOS were first detected, and then, after stripping (20 mn at RT in stripping buffer - catalog n°21063, Thermo Fisher Scientific, USA), the total amount of Akt or eNOS was measured to calculate phosphorylated protein/total protein ratio. Protein loading was normalized using HSP90 content, detected on the same membrane.

### Cell culture

Bovine aortic endothelial cells (BAEC, lot 1934, Cell applications Inc, CA, USA) at passages 5 to 8 were cultured at 37°C, 0,5% CO2 on 0.2% gelatin-coated 12-well plates or 35 mm Petri dishes using bovine cell growth medium (catalog n°B211, TEBU-BIO, France). Upon reaching confluence, cells were washed with warm HBSS without Ca^2+^ and Mg^2+^ and then treated for 1 hour with DMEM (catalog n°21885025, Thermo Fisher Scientific, MA, USA) containing 5% FBS or 5% plasma, obtained by centrifugation (2800 rpm, 20 minutes) of blood collected from the hepatic vein in a feeding state using heparinized tubes, from WT ND, WT HF, or Foz HF.

At the end of the treatment, cells were either exposed to sodium orthovanadate (1 mM) and collected for protein extraction to assess eNOS phosphorylation by western blot, or used to measure ROS generation using the fluorescent dyes CM-H2DCFDA (catalog n°C6827, Thermo Fisher Scientific, MA, USA).

To measure reactive oxygen species, cells were incubated with 20 µM CM-H2DCFDA in HBSS for 30 minutes at 37°C, protected from light. After washing twice with HBSS and incubating for 20 minutes in complete DMEM (see supplementary methods) without phenol red, the cells were trypsinized and collected in glass tubes. They were then washed twice by centrifugation (1,500 rpm, 5 minutes, RT) and re-suspended in HBSS containing DAPI at 0,5μM (catalog n°D9542, Sigma-Aldrich, Merck, Germany). Fluorescence was then measured using flow cytometry system (BD FACSCantoTM II, BD Biosciences, NJ, USA). H₂O₂ (catalog n° H1009, Sigma-Aldrich, Merck, Germany) at 15 µM for 1 min was used to induce ROS production as a positive control in each tube after the first measurement.

### Statistical analysis

Data are displayed as mean ± standard error of mean. Parametric data were analyzed using two-way or one-way analysis of variance with subsequent Bonferroni or Tukey’s multiple comparisons test, or unpaired t-test. For non-parametric data sets, statistical differences between groups were evaluated using Kruskal-Wallis test followed by Dunn’s multiple comparisons test or Mann-Whitney test. All analyses were performed with GraphPad Prism 9.2.0 (GraphPad Software, Inc., USA). Differences were considered statistically different with a *p*-value < 0,05. The significance levels are displayed as follows: *p<0.05, **p<0.01, ***p<0.001, ****p<0.0001. Data are depicted as mean ± standard error of mean. The utilized number of animals or vessel rings for each experiment is noted in the corresponding figure caption, with graphs displaying individual values.

## Results

### Foz mice develop MASH after 24 weeks on a high-fat diet

As previously reported^26,34,35^, the Foz genotype, when combined with a high-fat diet (Foz HF) for 24 weeks, led to severe obesity, with body weight reaching 70 grams (**Figure 1.B**). After a four-hour fast, glycemia did not differ significantly between the groups (**Figure 1.C**). However, we previously showed that mice with MASH exhibited elevated insulinemia at this stage, indicating severe insulin resistance^26^. The liver of Foz HF mice displayed severe panlobular steatosis, pronounced inflammation, and ballooning (**Figure 1.D**). In contrast, WT mice under normal diet (ND) mice had normal livers, while the livers of WT HF and Foz ND mice showed an intermediate phenotype, with moderate steatosis, mild inflammation, and inconsistent ballooning. In line with our previous findings, histological analysis of steatosis, ballooning, and inflammation showed that the NAFLD activity score (NAS) was highest in Foz HF (across all three parameters in each animal), intermediate in Foz ND anf WT HF, and absent in WT (**Figure 1.E**).

**Figure 1.**
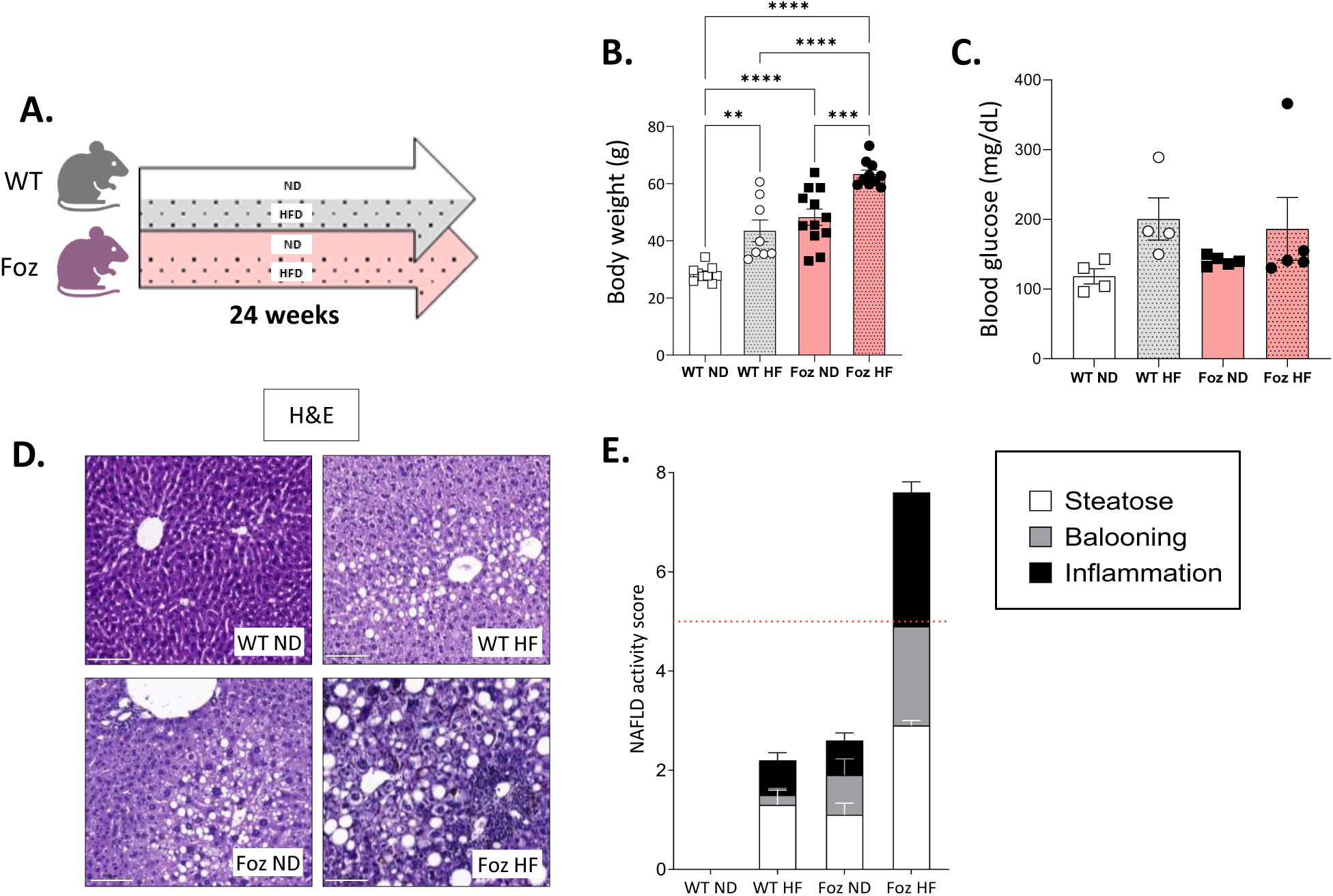
A. Experimental setup, B. Body weight and C. glycemia after 4h of fasting. D. Representative histological pictures of H&E staining; scale bar : 100μm. E. NAFLD acitivity score (NAS) based on evaluation of steatosis (white bars), ballooning (grey bars) and inflammation (black bars), threshold for MASH displayed as dashed red line; n= 4-12 mice per group ; statistics one-way ANOVA with Tukey’s multiple comparisons test (B) or Kruskal–Wallis with Dunn’s multiple comparisons test (C). Data are shown as mean +/- SEM, *p<0,05, **p<0,01, ***p<0,001, ****p<0,001.H&E:hematoxylinandeosin,HF:high-fat,ND:normaldiet,WT:wildtype

### Mice with MASH (Foz HF) but not those with HFD-induced steatosis have NO and EDHF(s)- dependent endothelial dysfunction

To assess endothelial function, we measured vasorelaxation in first order mesenteric arteries in all four groups of mice ranging from mice with healthy livers to those with MASH livers (**Figure 2.A**, **Supplemental Figure 1.A**). Vascular relaxation (expressed in % of maximal contraction) in response to increased concentrations of carbachol was evaluated. Phenylephrine (phe) or high KCl solution (50mM) were used to induce contraction (**Figure 2.B**). Isolated vessels from all four groups developed similar basal tension but the maximum contractile response to high KCl solution was larger in vessels from mice fed an HFD than in those on ND, irrespectively of genotype (**Supplemental Figure 1.B**). To avoid biased conclusions regarding relaxation capacity because of their different maximal contraction, we thereafter separately compared WT ND with Foz ND and WT HF with Foz HF. Total Relaxation (all pathways) of WT ND and Foz ND vessels was similar (**Supplemental Figure 1.C**). By contrast, total endothelium-dependent relaxation of phe pre-constricted vessels was less pronounced in vessels from Foz HF than in WT HF, although not significantly (**Figure 2.C**). In order to isolate the NO-dependent component of the relaxation, vessels were incubated in the presence of indomethacin (INDO), a COX-inhibitor, and contracted in the presence of high KCl solution, to block Prostacyclin (PGI2)-dependent relaxation and endothelium-derived hyperpolarization factors (EDHFs), respectively (**Figure 2.D**). In such conditions, vessels of Foz HF mice relaxed significantly less than those of WT HF mice (**Figure 2.E**) while vessels from WT ND and Foz ND showed no differences in their relaxation capacity (**Supplemental Figure 1.D**). In the same manner, L-NAME with high KCl was used to block eNOS and EDHF(s), respectively, or L-NAME with INDO to concomitantly inhibit both NO and PGI2 pathways (in this case, vessels were precontracted with Phe) (**Figure 2.D**). This way we showed that the PGI2- pathway contributes only marginally to relaxation in all groups (**Figure 2.F, Supplemental Figure 1.E**) while first order mesenteric arteries of Foz HF mice exhibited reduced EDHF(s)-dependent vessel relaxation compared to those of WH mice (**Figure 2.G**). This was not seen in vessels from mice fed the regular diet (**Supplemental Figure 1.F**). Taken together our analyses of the relaxation capacity of first order mesenteric arteries proved reduced NO- and EDHF(s)-dependent vasodilation in response to treatment with carbachol in Foz HF mice with MASH compared to their WT littermates on HFD with simple steatosis.

**Figure 2.**
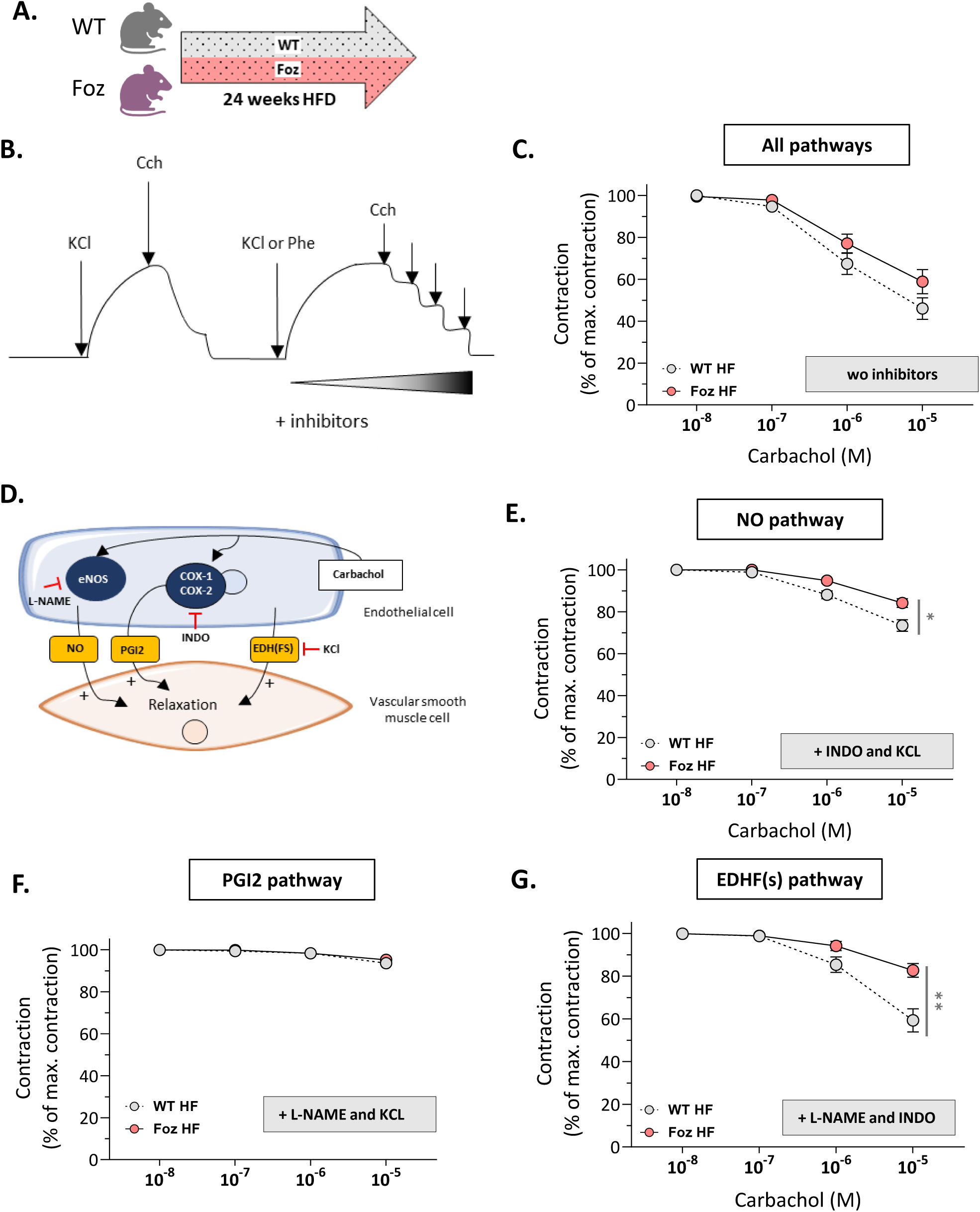
**A**. Experimental setup, **B.** Representative schematic of the experiment: vessels were contracted using phenylephrine (Phe 10^-7^M) and maximal relaxation was assessed with carbachol (Cch) at 10⁻⁵ M. Contraction was induced with either KCl (50 mM) or phenylephrine (10⁻^5^ M), and relaxation was assessed in response to increasing concentrations of carbachol**. C.** Total vasorelaxation in first order mesenteric arteries. **D.** Schematic representation of vasodilative pathways with corresponding inhibitors for selective suppression. **E**. NO- dependent relaxation with concomitant inhibition of alternative vasodilating PGI_2_- and EDHF-pathways (with INDO and KCL, respectively), **F.** PGI_2_-dependent relaxation examined by concomitant inhibition of alternative vasodilating NO- and EDHF-pathways (with L-NAME and KCL, respectively), **G.** EDHF(s)-dependent relaxation examined by concomitant inhibition of alternative vasodilating NO- and PGI_2_-pathways (with L-NAME and INDO, respectively) ; n= 20-31 rings per group ; statistics two-way ANOVA with Tukey’s multiple comparisons test (C,E,F and G). Data are shown as mean +/- SEM, *p<0,05, **p<0,01, ***p<0,001, ****p<0,001. Cch : carbachol, COX : cyclooxygenase, EDHF(s) : endothelium-derived hyperpolarizing factor(s), eNOS : endothelial NO synthase, HF : high-fat, INDO : indomethacin, L-NAME : L-N^G^-Nitro arginine methyl ester, PGI_2_ : prostaglandine I2, Phe : phenylephrine, NO : nitric oxide, ND : normal diet, WT : wild type.

### Blood pressure regulation, driven by the nitric oxide pathway and circadian rhythms, is disrupted in Foz mice on a high-fat diet

HFD-fed Foz or WT mice were treated with L-NAME (2g/L) in the drinking water for 7 days (**Figure 3.A**). Blood pressure was continuously recorded over 24 hours prior and following L-NAME treatment. At baseline, no significant differences were observed between the two groups with Foz and WT mice displaying similar mean systolic and diastolic blood pressure (SBP and DBP) (**Figure 3.B and D**). As expected, L-NAME induced an increase in SBP and DBP in both Foz and WT mice, with WT mice displaying significantly higher SBP at hours 4 and 20 of recording. This hypertensive response was attenuated in Foz mice (**Figure 3.B**). However, the increases in average 24-hour SBP and DBP induced by L-NAME treatment compared to the basal state were not statistically different between Foz and WT mice, likely due to high inter-individual variability (**Figure 3.C and E**). In line with the results presented in Figure 2, our observations support that NO–mediated blood pressure regulation is impaired in Foz mice with MASH.

**Figure 3.**
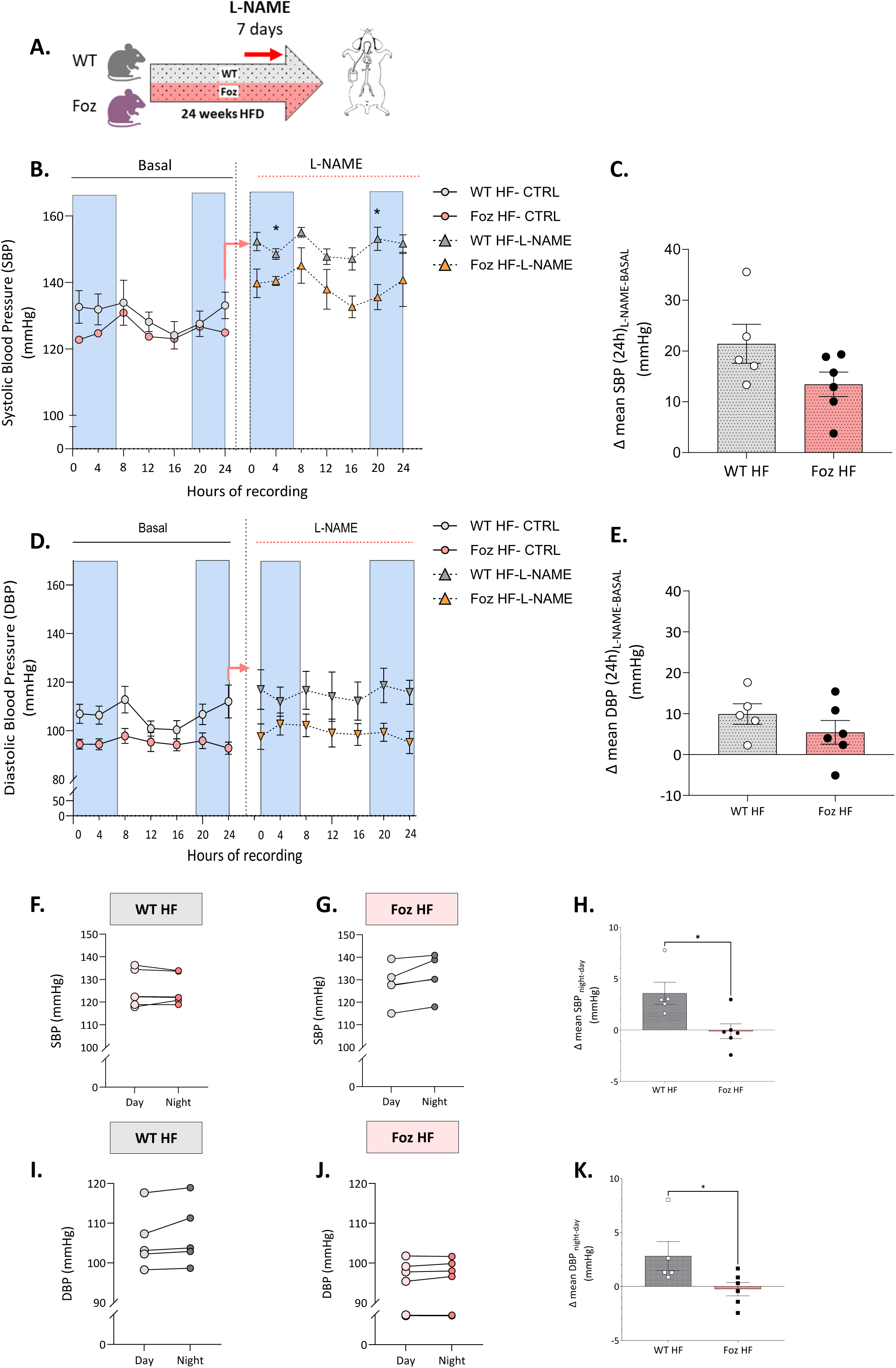
A. Experimental setup, **B.** Systolic blood pressure (SBP) recorded over 24 hours in the basal state or following L- NAME treatment. **C.** Difference in mean SBP between basal and L-NAME treatment. **D.** Diastolic blood pressure (DBP) recorded over 24 hours in the basal state or following L-NAME treatment **E.** Difference in mean DBP between basal and L-NAME treatment. **F, G.** Day–night variation in mean SBP in WT or Foz HF mice. **H.** Difference in mean SBP between day and night. **I, J.** Day–night variation in mean DBP in WT or Foz HF mice. **K.** Difference in mean DBP between day and night. n= 5-7 mice per group ; statistics two-way ANOVA followed by post hoc multiple comparisons with Bonferroni correction (B and D), unpaired two-tailed t-test (C) or Mann-Whitney test (F and I). Data are shown as mean +/- SEM, *p<0,05, **p<0,01, ***p<0,001, ****p<0,001. DBP : diastolic blood pressure ; L- NAME : L-NG-nitroarginine methyl ester ; SBP : systolic blood pressure; HF : high-fat; WT : wild type.

Under normal physiological conditions, blood pressure follows a circadian rhythm, characterized by a noticeable dip during the rest period (daytime). Interestingly, Foz mice exhibited a ’non- or reverse- dipper’ profile, showing a complete absence of nychthemeral variation in SBP and DBP compared to WT controls (**Figure 3.F, G, I and J**). Consequently, the difference in mean SBP and DBP between the active (night) and rest (day) periods was significantly reduced in Foz mice (**Figure 3.H and K**). Elevated blood pressure during the rest phase has been associated with adverse cardiovascular outcomes^36^ and in clinical practice, the day-to-night blood pressure ratio is a predictor of major cardiovascular events, even in normotensive individuals^37^.

### NO-control of vasorelaxation is impaired in an independent model of MASH

As an independent model of MASH, C57BL/6JRj were fed a western-type diet (40% kcal from fat, 40% kcal from sucrose and 0,5% of cholesterol) supplemented with 30% fructose in drinking water for 26 weeks. Mice on a normal diet (ND) were the healthy controls, those fed a regular HF with or without additional fructose (HF or HF+F) served as intermediate controls (**Supplemental Figure 2.A**). HF and HF+F mice showed a significant increase in body weight compared to the control group on a ND. Mice on the western diet with fructose (WD+F) exhibited an intermediate body weight (**Supplemental Figure 2.B**). This was accompanied by elevated fasting blood glucose levels in the HF, HF+F and WD+F groups compared to the control, although hyperglycemia appeared less pronounced in the WD+F group (**Supplemental Figure 2.C**). We measured an increased liver weight in WD+F associated with massive steatosis. HF and HF+F mice exhibit intermediate liver weight and moderate steatosis while ND mice had a healthy liver (**Supplemental Figure 2.D and 2.E**). Hepatic collagen content was significantly increased in the HF+F and more patently so in WD+F, as determined by the percentage of Sirius red positive area (**Supplemental Figure 2.E and 2.F**). Inflammation, assessed by F4/80 immunolabeling, revealed marked macrophage infiltration and prominent crown-like structures in the WD+F group. In contrast, HF and HF +F groups showed moderate activation of liver macrophages, while the ND group exhibited no signs of hepatic inflammation (**Supplemental Figure 2.G**).

The estimated NAFLD Activity Score (NAS), based on H&E staining, was high (around 6) in the intermediate groups HF and HF+F, and reached the maximum score of 8 in the WD+F group. In contrast, ND mice exhibited healthy liver histology with a NAS nearing 0 (**Supplemental Figure 2.H**). Based on these data, we conclude that C57BL/6JRj mice fed a WD+F diet represent a faithful model of severe, fibrosing MASH. Mice receiving a HFD, with or without fructose supplementation, develop MASH of intermediate severity without fibrosis, whereas ND-fed mice serve as appropriate controls with healthy liver.

In this independent model (**Figure 4.A**), we assessed vasodilation in superior mesenteric artery (expressed in % of maximal contraction) in response to increasing concentrations of carbachol. Contraction was induced using either phe or high KCl solution (20 mM). Total vascular relaxation, as well as relaxation mediated by PGI₂ alone or by both PGI₂ and NO pathways, did not differ between the four groups (**Figure 4.B, C and D**). To further investigate the role of NO pathway, we compared vessel relaxation before and after adding L-NAME to the organ bath in the myograph setup. As shown in **Figure 4.E**, L-NAME completely abolished carbachol-induced vasorelaxation in ND mice, indicating that the NO pathway plays a predominant role in vasorelaxation in these animals. In contrast, the response to L-NAME of vessels of WD+F animals with severe MASH was not significant regardless the carbachol concentration (two-way ANOVA; treatment factor p=0,1079) (**Figure 4.F**). These data suggest that the ability of the NO pathway to regulate vascular tone is impaired in an independent model of MASH-affected mice. In HF and HF+F animals, though to a lesser extent than in controls, L-NAME reduced the magnitude of vasorelaxation with p-value < 0,05 (**Supplemental figure 3.B and 3.C**).

**Figure 4.**
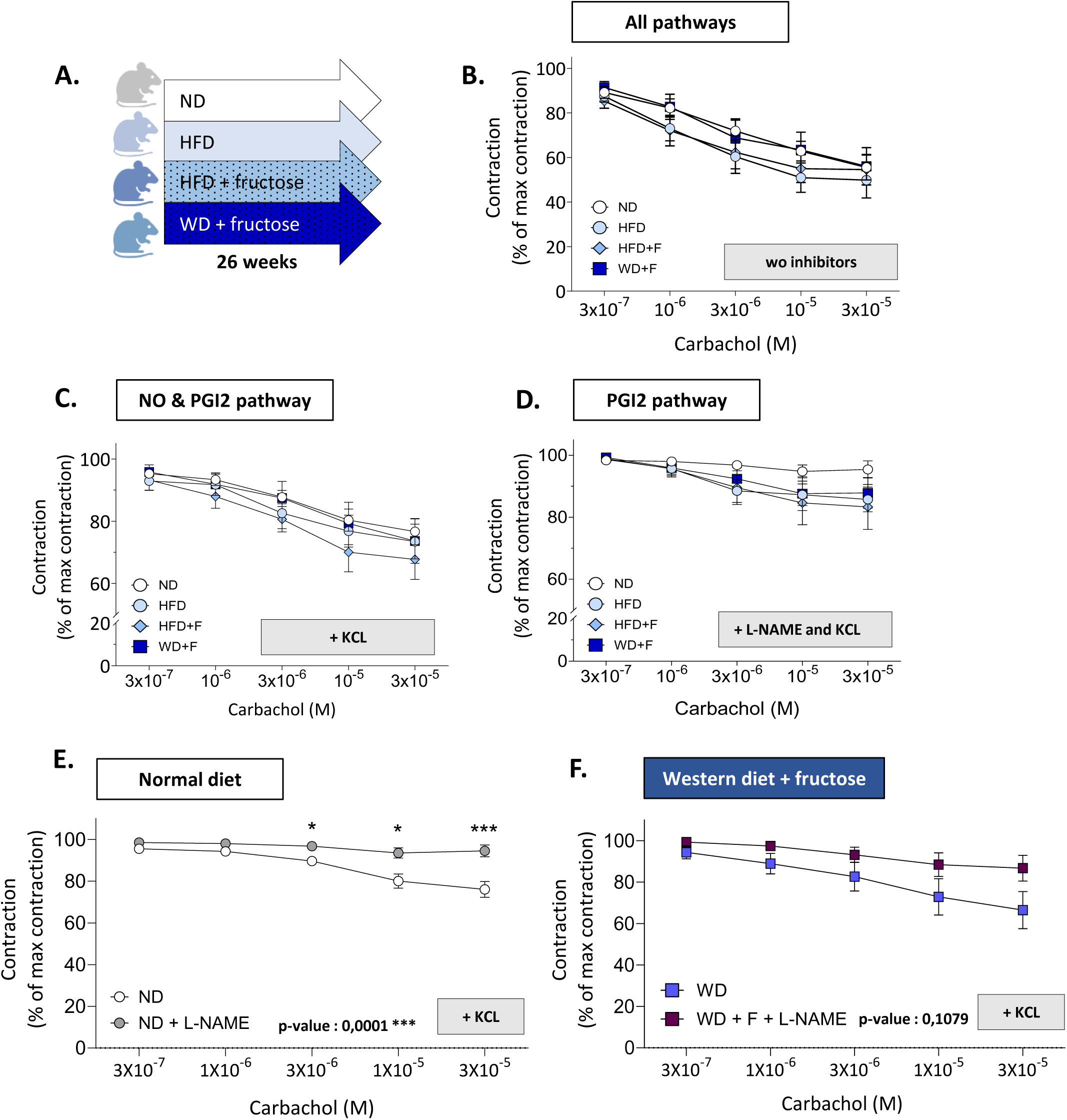
A. Experimental setup, **B.** Total vasorelaxation in superior mesenteric arteries. **C.** NO and PGI_2_-dependent relaxation examined by inhibition of vasodilating EDHF-pathways (with KCL). **D.** PGI_2_-dependent relaxation examined by concomitant inhibition of alternative vasodilating NO- and EDHF-pathways (with L-NAME and KCL, respectively) **E**. Vasorelaxation in superior mesenteric of ND (**E**), and WD+F (**F**) induced by KCL (EDHF pathway inhibitor) with or without L-NAME treatment. The reported p-value corresponds to the effect of the treatment factor (L-NAME) as determined by two-way ANOVA (E,F) ; n= 12-16 rings per group ; statistics two-way ANOVA followed by post hoc multiple comparisons with Bonferroni correction. Data are shown as mean +/- SEM, *p<0,05, **p<0,01, ***p<0,001, ****p<0,001. Cch : carbachol, EDHF(s) : endothelium-derived hyperpolarizing factor(s), HF : high-fat, INDO : indomethacin, L-NAME : L-N^G^-Nitro arginine methyl ester, PGI_2_ : prostaglandine I2, Phe : phenylephrine, NO : nitric oxide, ND : normal diet, WT : wild type.

### Impaired NO-dependent vasorelaxation is associated with reduced *eNOS* in mice with MASH

To identify the cause of loss of vasoactive response to NO, we further evaluated the NO signaling pathway in the aorta of Foz as well as C57BL6/JRj mice (**Figure 5.A, Supplemental figure 3.A**). Compared to their dysmetabolic controls (WT HF) the expression of *eNOS*, the main producer of NO in vascular endothelial cells, was lower in the aorta of Foz mice (**Figure 5.B**). No difference was observed in the expression of *Cav1* and *Arg1*—two negative regulators of NO production—between HF-fed Foz mice and WT (**Figure 5.B**). Phosphorylation of eNOS at serine ^1177^ (p^Ser1177^ eNOS) is known to enhance eNOS activity and subsequently increase NO production. Western blot analysis revealed reduced eNOS activation, as evidenced by decreased p^Ser117^, in the aortas of HF-fed Foz mice (**Figure 5.C and D**). Thus, the data support a decreased production of NO in Foz HF aortas, relative to WT HF. At variance, the observations were not replicated in the second model (**Supplemental figure 3.D, E and F**).

**Figure 5.**
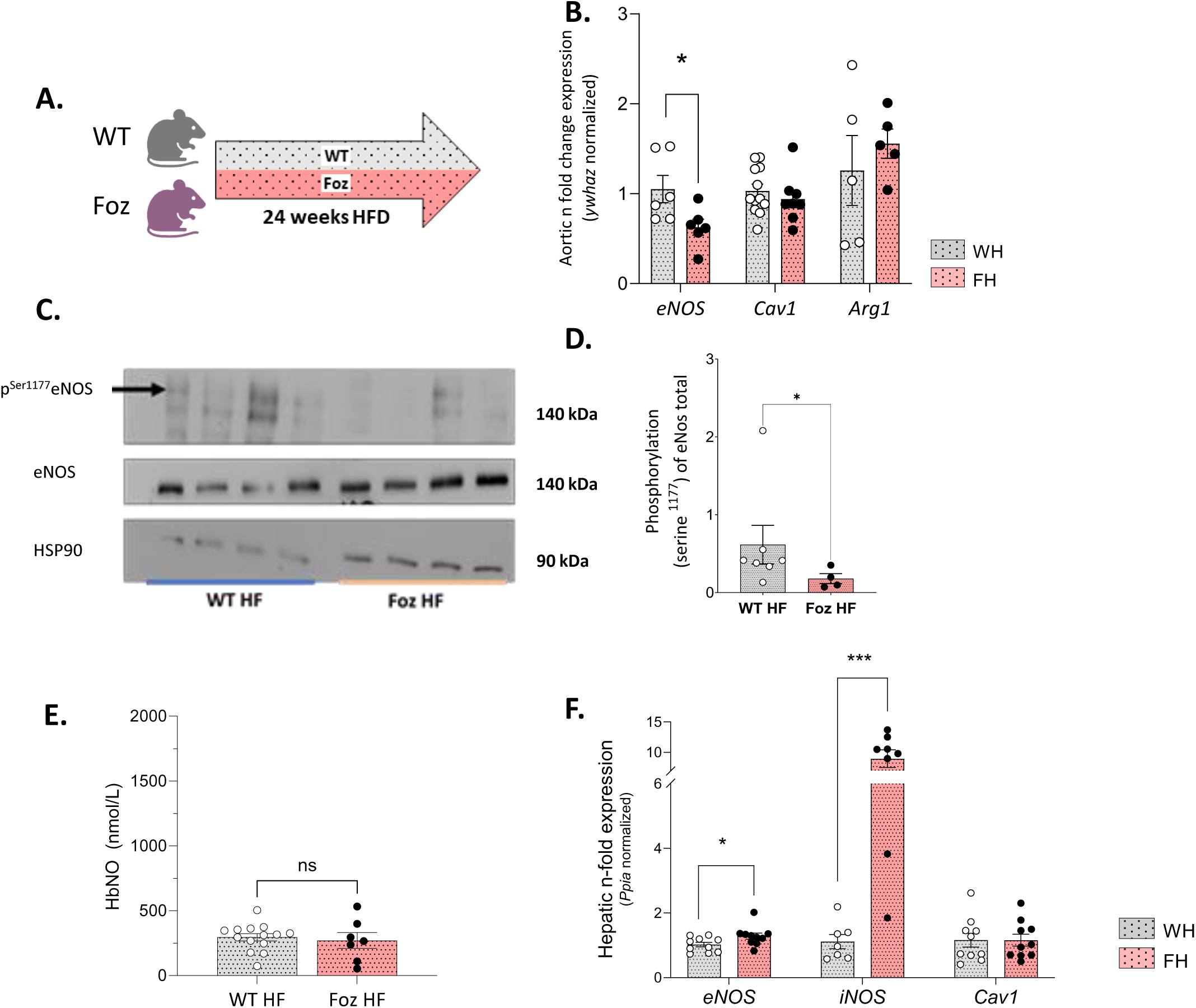
A. Experimental setup **B.** Aortic gene expression levels normalized with *Ywhaz, n*-fold expression using WT HF as reference **C.** Detection of phosphorylated Serine^1177^ form of eNOS, total eNOS and HSP90 by western blot in aorta of HF-fed Foz and WT mice. **D.** Quantification of phosphorylated Serine^1177^ form of eNOS on total eNOS. HSP90 was used as a loading control. **E.** HbNO level in whole blood. **F.** Hepatic gene expression levels normalized with *ppia, n*-fold expression using WT HF as reference; n=4-10 mice per group ; statistics : Mann-Whitney test (D,F); unpaired two-tailed t-test (B,E). Data are shown as mean +/- SEM, *p<0,05, **p<0,01, ***p<0,001, ****p<0,001. Cav1 : caveolin1, eNOS : endothelial nitric oxide synthase, HbNO : nitrosylated hemoglobin, HF : high-fat, iNOS: inducible nitric oxide synthase, Ppia : peptidylprolyl isomerase A, WT : wild type, ywhaz : 14-3-3-zeta.

We then measured blood nitrosylated hemoglobin (HbNO), a surrogate for *in vivo* NO production. Despite Foz mice having a markedly low HbNO levels compared to healthy controls under ND (**Supplemental Figure 4.C**), HF-fed Foz and WT mice exhibited similar HbNO concentrations (**Figure 5.E**) indicating the HFD reduces NO availability in WT but not in Foz mice. This observation suggests the presence of an alternative NO source in HF-fed Foz mice with MASH. We therefore explored hepatic NO production. Gene expression analysis in the liver revealed a 17-fold upregulation of inducible nitric oxide synthase (iNOS), likely driven by hepatic inflammation, in Foz mice compared to dysmetabolic controls (WT HF) (**Figure 5.F**). In contrast, Foz mice on ND with fatty liver exhibited only a modest, non-significant increase in *iNOS* expression relative to healthy controls (**Supplemental Figure 4.D)**. Additionally, hepatic *eNOS* expression was also slightly elevated in Foz HF compared to HF-fed WT mice, while expression levels of *Cav1* remained unchanged (**Figure 5.F**). Of note, we also observed a marked increase in *iNOS* expression in C57BL/6JRj WD+F mice compared to ND, supporting the notion that MASH is associated with a substantial release of NO driven by liver inflammation (**Supplemental figure 3.G**).

Taken together, our findings demonstrate dysregulation of aortic eNOS in MASH-afflicted Foz mice, both at the transcriptional and post-translational levels, compared to their HF-fed WT littermates. However, these alterations, known to impair NO production, were not reflected in circulating HbNO levels. In this context, the markedly increased hepatic *iNOS* expression suggests that the inflamed liver may act as an alternative source of excessive NO in MASH. Consequently, in such a context, HbNO does not accurately reflect NO production within the vasculature.

### In MASH, circulating factors inhibit serine^1177^ eNOS phosphorylation *ex vivo*

To investigate potential hepatic drivers that may inhibit eNOS activation in the context of MASH, we treated BAECs for 1 hour with 5% plasma from WT ND mice as a healthy control, WT HF mice as a dysmetabolic control, and Foz HF mice as a model of MASH. After treatment, proteins were collected, and we assessed eNOS activation by measuring p^Ser1177^ eNOS using western blot. We observed a strong trend toward decreased p^Ser1177^ eNOS following treatment with plasma from Foz HF mice compared to controls, although the difference was not statistically significant after quantification (p-value: 0.059) (**Figure 6.A and B**).

**Figure 6.**
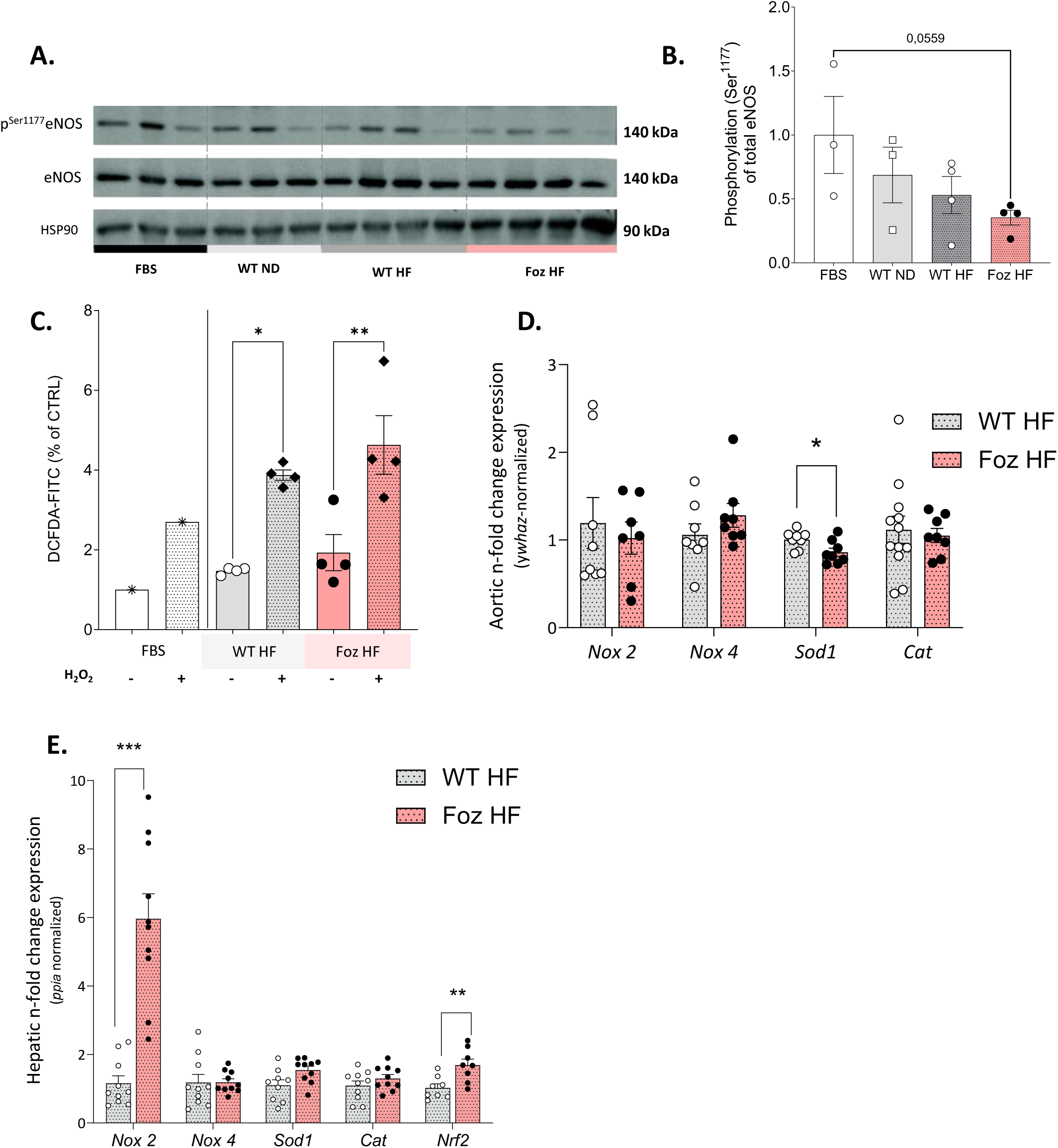
A. Detection of phosphorylated Serine^1177^ form of eNOS, total eNOS and HSP90 by western blot in BAEC following FBS or plasma treatments. **B.** Quantification of phosphorylated Serine^1177^ form of eNOS on total eNOS. HSP90 was used as a loading control. **C.** DCFDA fluorescence in BAEC cells following plasma treatments and H_2_O_2_ treatment, results as expressed in % of control (FBS treatment). **D.** Aortic gene expression levels normalized with *ywhaz, n*- fold expression using WT HF as reference, **D.** Aortic gene expression levels normalized with *Ppia, n*-fold expression using WT HF as reference. n=4-11 mice per group ; statistics : unpaired two-tailed t-test (D,E), one-way ANOVA with Tukey’s multiple comparisons test (B,C). Data are shown as mean +/- SEM, *p<0,05, **p<0,01, ***p<0,001, ****p<0,001. Cat : catalase, FBS: fetal bovine serum, HF : high-fat, Nox 2/4 : NADPH oxidase 2/4, Nrf2 : nuclear factor (erythroid-derived 2)-like 2, Sod1 : superoxide dismutase 1, Ppia : peptidylprolyl isomerase A, WT : wild type, ywhaz : 14-3-3-zeta.

We used the fluorescent marker DCFDA to determine whether Foz HF plasma treatment induces oxidative stress in BAECs, potentially leading to inhibition of eNOS. H2O2 was employed as a positive control to induce the production of reactive oxygen species (ROS). As shown in **Figure 6.C**, H2O2 treatment significantly increased DCFDA fluorescence in the cells. However, no significant differences were observed between cells exposed to FBS (controls) and cells treated with plasma from either HF- fed WT or Foz mice. Consistent with these results, no significant changes were detected in the aortic expression of genes involved in redox balance, except for a slight decrease in superoxide dismutase 1 (*Sod1*) expression in the aorta of Foz HF mice (**Figure 6.D**).

Thus, we have demonstrated that plasma-derived factors can inhibit eNOS through Serine^1177^ dephosphorylation in endothelial cells in an oxidative stress independent manner. Moreover, although we observed increased expression of oxidative stress markers (*Nox2* and *Nrf2*) in the liver of Foz mice compared to WT (**Figure 6.E**), a similar increase was not detected in the vasculature. Therefore, despite the well-established role of oxidative stress in endothelial dysfunction, it does not appear to be a driving factor in vascular dysfunction in the context of MASH.

### Hepatic and plasma ADMA levels are elevated in two independent MASH Models

ADMA, a methylated derivative of L-arginine, is a key regulator of NO production. It is synthesized throughout the body by protein arginine N-methyltransferases (PRMTs) and predominantly metabolized in the liver and kidney by dimethylarginine dimethylaminohydrolases (DDAHs). The isoform *Ddah2* is mainly expressed in vascular endothelial cells, whereas *Ddah1* is the major hepatic isoform. In two independent models of MASH (Foz mice on HF and C57BL/6JRj mice fed WD supplemented with fructose), we observed elevated ADMA levels in both plasma and liver compared to their respective controls (**Figure 7.B, C, F and G**). In the Foz model, plasma ADMA levels gradually increased across groups in parallel to the worsening severity of liver injury (**Supplemental Figure 5B**). A similar pattern was observed for plasma and hepatic ADMA levels in C57BL/6JRj mice (**Figure 7.F and 7.G**), suggesting a potential link between the severity of liver disease and ADMA accumulation in liver and bloodstream.

**Figure 7.**
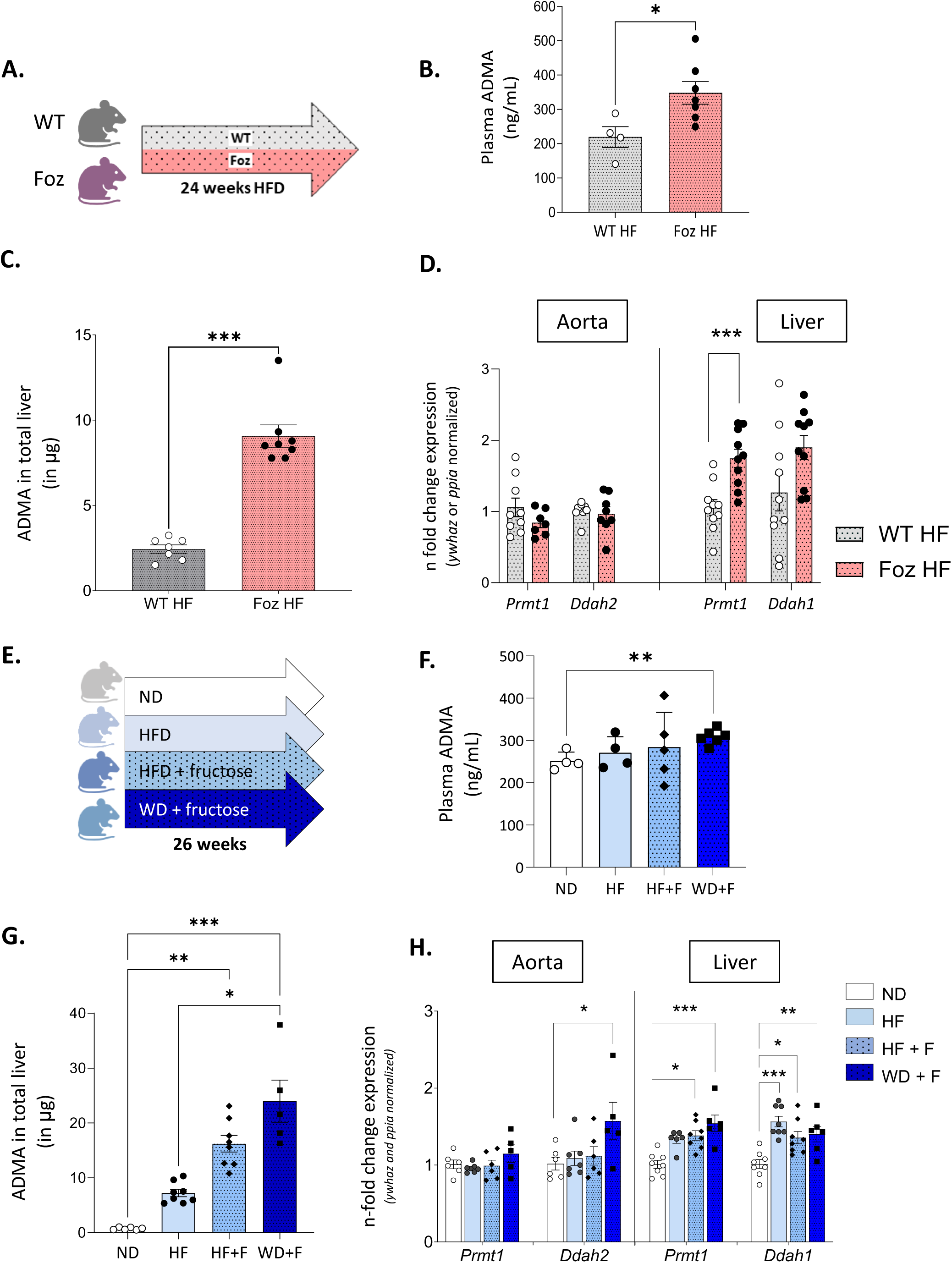
A. Experimental setup. **B.** Plasma ADMA levels in ng/ml of plasma, **C**. ADMA content in total liver (in μg) . **D**. Aortic and hepatic *Prmt1 and Ddha1/2* gene expression levels normalized with *ywhaz* and, *Ppia* respectively, n-fold mRNA expression calculated using WT HF as reference. **E.** Experimental setup. **F.** Plasma ADMA levels in ng/ml of plasma, **G**. ADMA content in total liver (in μg) . **H**. Aortic and hepatic *Prmt1 and Ddha1/2* gene expression levels normalized with *ywhaz* and *Ppia, r*espectively, n-fold mRNA expression calculated using WT HF as reference. n=4- 10 mice per group ; statistics : unpaired t-test (**B,D**), unpaired Mann-Whitney test (**C**), one-way ANOVA with Tukey’s multiple comparisons test (**F,G**) or Kruskal–Wallis with Dunn’s multiple comparisons test (**H**). Data are shown as mean +/- SEM, *p<0,05, **p<0,01, ***p<0,001, ****p<0,001. ADMA : asymmetric dimethylarginine, Ddah1/2 : dimethylarginine dimethylaminohydrolase 1/2 F : fructose, HF : high-fat, ND : normal diet, Ppia : peptidylprolyl isomerase A, Prmt1: protein arginine methyltransferase 1, WD : western diet, WT : wild type, ywhaz : 14-3-3-zeta.

Furthermore, hepatic *Prmt1* expression was significantly upregulated in both Foz HF and C57BL/6JRj WD+F mice compared to their respective controls (**Figure 7.D and 7.H**). In the C57BL/6JRj model, but less so in Foz with intermediate liver disease (**Supplemental Figure 5.B, C and D**), *Prmt1* expression closely mirrored hepatic ADMA levels and increased with the severity of liver pathology. In the aorta however, *Prmt1* expression did not differ significantly between healthy and MASH-afflicted mice (**Figure 7.D and 7.H**), further supporting the hypothesis that hepatic (and not locally produced) ADMA may drive eNOS inhibition. In HF-fed Foz mice, no significant changes were observed in *Ddah2* expression in the aorta or *Ddah1* expression in the liver compared to dysmetabolic controls WT HF (**Figure 7.D**). In C57BL/6JRj mice, *Ddah2 and Ddah1* expression was elevated in the aorta and the liver of WD+F relative to healthy controls (**Figure 7.H**). Collectively, these findings suggest that in MASLD/MASH, upregulation of *Prmt1* may drive hepatic ADMA accumulation, with subsequent release into the bloodstream. Circulating ADMA could then impair NO production in endothelial cells, thereby contributing to endothelial dysfunction.

## Discussion

Our study provides compelling evidence that endothelial dysfunction is a key feature associated with MASH in two independent murine models. Whether in diet-induced or in genetically predisposed models, we demonstrated that NOS/NO-dependent endothelial impairment is not merely a coincidental occurrence but may represent a pathophysiological link between metabolic hepatic stress and systemic endothelial alterations. Consistent with previous reports indicating vascular dysfunction in MASLD patients^17–19^, our findings extend the paradigm to MASH and provide mechanistic insights as we identified ADMA as a potential MASH-driven contributor to vascular impairment.

We recently demonstrated that MASH in HF-fed Foz mice promotes cardiac abnormalities, namely cardiac hypertrophy, fetal gene reprogramming, and myocardial fibrosis, thereby priming the heart toward cardiac dysfunction^26^. We extend these findings by showing impaired NO-dependent arterial vasorelaxation in HF-fed Foz mice, associated with reduced eNOS expression and activation. NO-related vascular alterations were also observed (though to a lesser extend) in an independent model as attested by the reduced dependence of arterial relaxation response on the NOS/NO pathway in C57BL/6JRj mice with MASH. The observed difference between models may, in part, be attributed to the distinct vascular segments used for functional assessment: vasorelaxation was evaluated in Foz in first-order mesenteric arteries (FMA), resistance-sized arteries with a prominent role in regulating vascular tone^38^ while in the superior mesenteric artery (SMA), a conduit vessel, in the C57BL/6JRj model. Nevertheless, we observed in the aorta of the later increased expression of cyclooxygenase-1 (COX-1), a key enzyme in PGI₂ synthesis, consistent with previous observations in atherosclerosis models^39^. The upregulation of the PGI₂ pathway may mask a reduction in NO-mediated vasorelaxation when the 2 pathways are evaluated together in the wire myograph. We also observed distinct patterns of eNOS expression and activation between the two models as eNOS expression was increased in C57BL/6JRj but not in Foz mice. This may nevertheless indicate a dysfunctional status of the vessels. For instance, Katakam et al. reported impaired insulin-induced vasodilation in the coronary arteries of obese Zucker rats compared to lean controls, alongside eNOS upregulation, probably reflecting a compensatory response by the endothelium, to counteract NO depletion and support vascular function^40^.

To test whether circulating factors contribute to the vascular phenotype, we incubated endothelial cells with plasma from mice with MASH. We observed reduced eNOS activation in the absence of significant oxidative stress. This supports that a circulating factor has disrupted NO signaling and subsequently decreased NO production through a mechanism independent of oxidative stress, although we did not test for the potential role of blood cells in MASH as activated platelets, which may promote oxidative stress and subsequent endothelial dysfunction^41^.

Interestingly, despite differences in the extent of endothelial dysfunction, the two murine models exhibited elevated plasma and hepatic ADMA levels. ADMA is an endogenous inhibitor of NO synthesis: It competes with L-arginine for transport and eNOS binding site, and reduces eNOS activation by lowering Ser¹¹⁷⁷ phosphorylation^42,43^. Literature consistently shows that elevated ADMA levels correlate with both the severity and the risk of cardiovascular disease^44,45^ and serve as an early marker of endothelial dysfunction^46^. In human volunteers, the administration of a pathophysiological bolus dose of ADMA significant increased blood pressure and systemic vascular resistance^47^. ADMA could therefore play an active role in endothelial dysfunction, beyond its established role as a biomarker. By contrast, our data show that Hb-NO may not be a reliable marker of endothelial dysfunction in inflammatory conditions such as MASH. Indeed, *iNOS* upregulation in these contexts might mask reduced NO production by vascular endothelial cells, an effect previously observed in sepsis patients^48^.

Building on these considerations, we hypothesize that elevated plasma levels of ADMA in MASH contribute to reduce NO production in vascular endothelium, ultimately leading to endothelial dysfunction. It could be argued that, because intracellular ADMA concentrations are 10 times higher in the cells (10 µM) than in the plasma (1 µM), elevated plasma ADMA alone may not be sufficient to significantly compete with endothelial NO synthesis^49^. Nevertheless, studies have shown that exposure of human endothelial cells to ADMA at 0,1μM is already sufficient to disrupt NO production^50^. Furthermore, because ADMA and arginine share the same cellular transporter (e.g., CAT-1)^51^, elevated plasma ADMA could competitively limit arginine uptake into endothelial cells, thereby impairing NO production even without directly affecting eNOS. Systemic reduction in arginine bioavailability due to MASH condition, also reflected by lower arginine/ornithine ratio in the liver, may additionally impair endothelial NO production independently of ADMA (**Supplemental figure 6**.) .

High plasma ADMA levels have been reported in patients with MASLD/MASH^52,53^, but the mechanisms linking liver disease with the increased production of eNOS inhibitor remain unclear. ADMA is generated during the degradation of proteins methylated on arginine residues. Protein arginine methyltransferases (PRMTs) catalyze the monomethylation and asymmetric dimethylation of these residues, with PRMT1 being the predominant isoform responsible for ADMA formation. Dimethylarginine dimethylaminohydrolases (DDAH1 and DDAH2) transform ADMA to citrulline and dimethylamine. In mice, DDAH1 or DDAH2 deficiency elevate plasma ADMA, causing hypertension and impaired vasoreactivity^54^, while *ddah1* overexpression alleviates hypertension^55^. Prior studies suggest that low DDAH1 activity, caused by oxidative stress and lipid overload, may promote ADMA accumulation in MASH-associated endothelial dysfunction^24^. The other way round, DDAH1 up- regulation has also been reported to protect against high-fat-diet-induced hepatic steatosis^56,57^. We found no systematic change in *Ddah1* expression in our models but did not measure its activity. Conversely, hepatic *Prmt1* expression was consistently upregulated. Metabolic disorders are associated with PRMT1 dysregulation^58^ and some studies have suggested that insulin resistance drives ADMA accumulation, through *Prmt1* overexpression^59,60^. Yet, in our C57BL/6JRj model, elevated plasma ADMA and hepatic *Prrmt1* mRNA levels were observed despite insulin sensitivity (data not shown). PRMT1 may contribute to MASLD/MASH though this view remains controversial. Some studies support that induction of PRMT1 is a compensatory mechanism to promote fatty acid oxidation via its action on PGC-1α^61^, while others propose that PRMT1, as it drives lipogenesis, contribute to steatosis^62,63^. This warrants investigation, as PRMT inhibition may not only counteract MASLD but also reduce ADMA levels and associated cardiovascular complications^26,64^ potentially positioning it as a therapeutic target.

Additionally, although MASH mice exhibit normal average blood pressure, altered nychthemeral (day- night) variations in SBP and DBP have been observed. Such a disruption in circadian blood pressure rhythms implies dysautonomy and is recognized as an important predictor of adverse cardiovascular outcomes ^65,66^. MASH had a strong effect on circadian liver transcriptome rhythms^67,68^. Whether and how the perturbed circadian metabolism in the liver affects autonomic function adversely and thus cardiovascular prognosis remains to be explored.

Several limitations to our study should be acknowledged. First, while murine models provide critical insights, they do not fully recapitulate the complexity of the human disease nor the cumulation of genetic, life-style and environmental factors. Second, in both mouse models studied, vascular abnormalities occurred in the absence of overt atherosclerosis (data not shown), confirming that endothelial dysfunction can precede or occur independently of plaque formation in metabolic liver disease. Mice are generally less susceptible to atherosclerosis than humans, largely due to their favorable lipoprotein profile - high HDL and low LDL levels. Consequently, only highly atherogenic diets, such as the Paigen diet (1% cholesterol and cholic acid), reliably induce atherosclerotic lesions after long exposure^69^. Also, the NOD genetic background of Foz mice may confer immune-dependent protection against plaque formation^70^. Finally, we only used male mice in our experiments. This choice is supported by the fact female mice are protected against ectopic fat accumulation and are consequently more resistant to MASH than males^71^. In addition, as estrogens and sex chromosome-linked genes promote NO production, females are also protected from endothelial dysfunction and its pathological consequences^72–74^.

In conclusion, our study demonstrates that NO-dependent endothelial dysfunction is a consistent feature of MASH in two distinct mouse models. We identified ADMA as a key liver-derived mediator of endothelial dysfunction in animals with MASH, a proposal that however does not exclude the contribution of other factors, circulating or not, such as miRNA or TMAO, to participate in the cardiovascular consequences of MASH^75–78^. The PRMT1/ADMA/NO axis appears to link hepatic pathology to vascular impairment independently of oxidative stress. Model-specific differences suggest the involvement of multiple, stage-dependent mechanisms contributing to endothelial dysfunction. These findings position ADMA as a potential biomarker and therapeutic target for MASH-associated cardiovascular risk, underscoring the need for further investigation into the role of PRMT1 in hepatic and circulating ADMA accumulation. From a translational perspective, our results suggest that endothelial dysfunction should be considered as a critical obligatory target in MASH patients to mitigate the risk of severe cardiovascular morbi- mortality.

## Authors contributions

Conceptualization : C.D and I.L ; methodology and data interpretation: C.D, I.L, J.L, S.B ; investigation : H.E (telemetry), J.L, K.S (LC-MS analyses), M.H, S.A (Flow cytometry protocol optimization), S.B and Z.B ; resources : C.D, I.L and O.F ; data curation : H.E, J.L, K.S, S.B ; writing – original draft : J.L; writing – review & editing : C.D and I.L ; visualization : J.L and S.B ; supervision : C.D and I.L ; funding acquisition : C.D and I.L. All authors approved of the final version of the manuscript.

The authors declare no conflict of interest.

## Funding

This work was supported by funding from the Fonds de la Recherche Scientifique (FNRS) with the project PDR THEMA-CARDIO: “CVD in NASH” P.C006.22 (to I.L. and C.D.). C.D. is a senior research associate with the FNRS, Belgium.

## Acknowledgements

We thank Faïza Amjahed, Natacha Feza-Bingi, Corinne Picalausa (GAEN/IREC/UCLouvain, Brussels, Belgium), Delphine de Mulder (FATH/IREC/UCLouvain, Brussels, Belgium) and Rachid El Kaddouri (ANIM/IREC/UCLouvain) for technical support. We thank Caroline Bouzin (2IP/IREC/UCLouvain, Brussels, Belgium) and Davide Brusa (Cytoflux/IREC/UCLouvain, Brussels, Belgium) for their expert assistance with image analysis and flow cytometry experiments.

## Declaration of Generative AI and AI-assisted technologies in the writing process

During the preparation of this work, the author(s) used the generative AI tool ChatGPT to enhance the manuscript’s readability and language. After using this tool/service, the author(s) reviewed and edited the content as needed and take(s) full responsibility for the content of the publication

## Notes

### Competing Interest Statement

The authors have declared no competing interest.

