## Supplements for "Hepatic ADMA–PRMT1 axis regulation is associated with NO-dependent endothelial dysfunction in MASH"

Table of contents

Supplementary methods............................................................................................ 2

Supplementary figures.............................................................................................. 3

### Supplementary methods

**Table 1. Primers used for gene expression analysis**

| Gene | Forward | Reverse |
|---|---|---|
| *Arg1* | ACAAGACAGGGCTCCTTTCAG | TGAGTTCCGAAGCAAGCCAA |
| *Arg2* | TACCATTATCGGTCACGCCC | GAGGCTCCACATCTCTCAGG |
| *Cat* | AGAGGAAACGCCTGTGTGAG | GCGTGTAGGTGTGAATTGCG |
| *Cav-1* | GACCCCAAGCATCTCAACGA | AGATGCCGTCGAAACTGTGT |
| *Ddah1* | GGTGGGGACGTCCTATTCAC | GAGGGCCTTCTGTGCAGATT |
| *Ddah2* | TTGCCTCAGTTTACCTCCTAGT | GGCTCTGGACCTGGCTAAAG |
| *eNOS* | TCTCACCTACTTCCTGGACATC | AGCTGAACCACTTCCATTCTTC |
| *iNOS* | GAAGGGGACGAACTCAGTGG | GTGGCTCCCATGTTGCATTG |
| *Ppia* | GGTGGAGAGCACCAAGACAGA | GCCGGAAGTCGACAATGATG |
| *Prmt1* | CACCCTCACATACCGCAACT | GCAAACATGCAGAGGATGCC |
| *Nox2* | CTGAAGGGGGCCTGTATGTG | CCAAACTCTCCGCAGTCTGT |
| *Nox4* | CCCTCCTGGCTGCATTAGTC | CGGTAAAGTCTCTCCGCACA |
| *Nrf2* | CTGCCATCAGTCAGTCACTCTC | CTCCGTAAATGGAAGACTCCAC |
| *Sod1* | GGGTTCCACGTCCATCAGTAT | GTACGGCCAATGATGGAATGC |
| *Ywhaz* | AGGCAGAGCGATATGATGAC | AATACTTGAGACGACCCTCC |

**Table 2. Krebs-Henseleit solution formulation**

| Component | Krebs-Henseleit buffer | High KCl-buffer (50 mM) |
|---|---|---|
| NaCl | 120.0 mM | 76.0 mM |
| KCl | 5.9 mM | 50.0 mM |
| $NaHCO_3$ | 25.0 mM | 25.0 mM |
| $NaH_2PO_4$ | 1.2 mM | 1.2 mM |
| $MgCl_2$ | 1.2 mM | 1.2 mM |
| $CaCl_2$ | 2.5 mM | 2.5 mM |
| glucose | 11.5 mM | 11.5 mM |

**DMEM without red phenol formulation**

DMEM (catalog n°D5030, Sigma-Aldrich, Merck, Germany)

5.5mM glucose (catalog n°G8270, Sigma-Aldrich, Merck, Germany)

2mM Glutamax  (catalog n°35050038 , Thermo Fisher Scientific, MA, USA)

10%FBS (catalog n°F7524, Sigma-Aldrich, Merck, Germany), heat inactivated (56°C, 30mn)

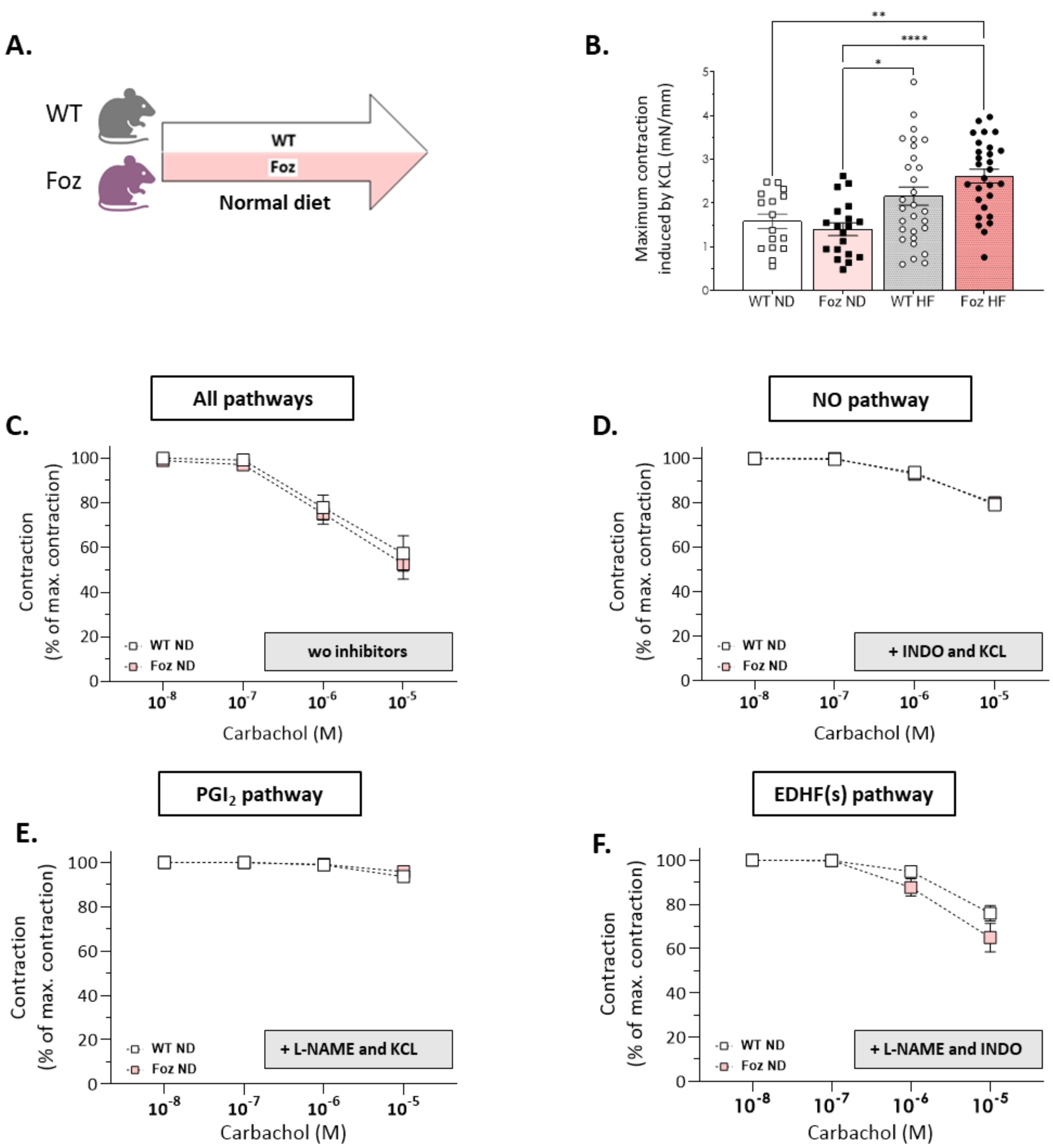


**Supplemental Figure 1. A.** Experimental setup, **B.** Maximum contraction (mN/mm) induced by KCL (50mM), C. Total vasorelaxation in first order mesenteric arteries. **D.** NO-dependent relaxation with concomitant inhibition of alternative vasodilating $PGI_2$- and EDHF-pathways (with INDO and KCL, respectively), **E.** $PGI_2$-dependent relaxation examined by concomitant inhibition of alternative vasodilating NO- and EDHF-pathways (with L-NAME and INDO, respectively) **F.** EDHF(s)-dependent relaxation examined by concomitant inhibition of alternative vasodilating NO- and $PGI_2$-pathways (with L-NAME and KCL, respectively) ; n= 16-32 rings per group ; Data are shown as mean +/- SEM, *p<0,05, **p<0,01, ***p<0,001, ****p<0,001. Statistics one-way ANOVA following Tukey's multiple comparisons test (**B**) or two-way ANOVA following Tukey's multiple comparisons test (**C, D, E** and **F**). EDHF(s) : endothelium-derived hyperpolarizing factor(s), HF : high-fat, INDO : indomethacin, L-NAME : L-$N^G$-Nitro arginine methyl ester, $PGI_2$ : prostaglandine I2, NO : nitric oxide, ND : normal diet, WT : wild type.

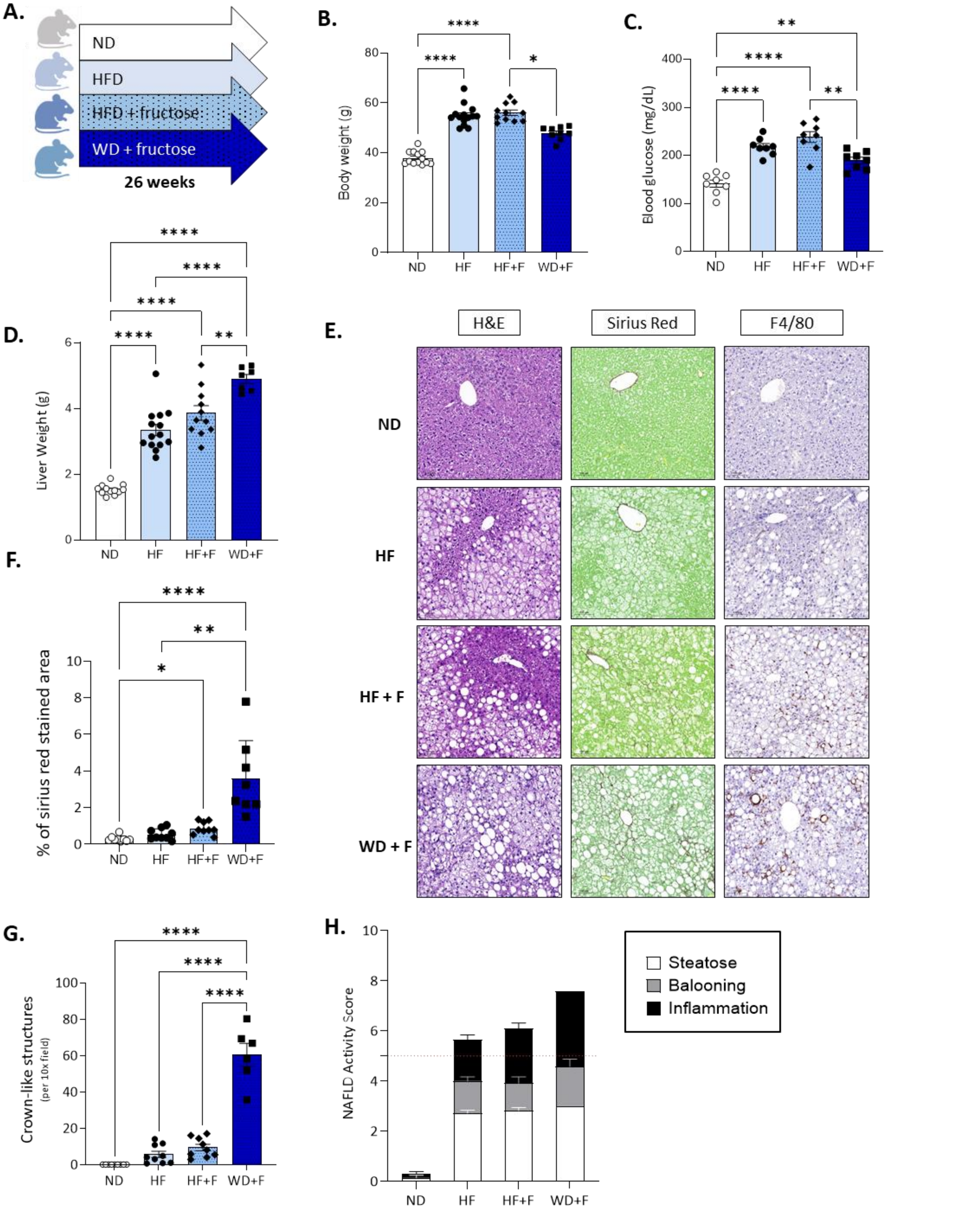


**Supplemental Figure 2.A.** Experimental setup, **B.** Body weight and **C.** glycemia after 4h of fasting. **D.** Liver weight. **E.** Representative histological pictures of H&E, Sirius red and F4/80 staining; scale bar : 100µm. **F.** Quantification of hepatic collagen content in % of total area. **G.** **N**umber of crown-like structures per 10× field of view (using aforementioned F4/80-stained sections); . **H.** NAFLD acitivity score (NAS) based on evaluation of steatosis (white bars), ballooning (grey bars) and inflammation (black bars), threshold for MASH displayed as dashed red line; n= 7-9 mice per group ; statistics one-way ANOVA with Tukey's multiple comparisons test (**B,D** and **G**) or Kruskal–Wallis with Dunn's multiple comparisons test (**A, F**). Data are shown as mean +/- SEM, *p<0,05, **p<0,01, ***p<0,001, ****p<0,001. F : fructose, HF : high-fat, ND : normal diet, WD : western diet.

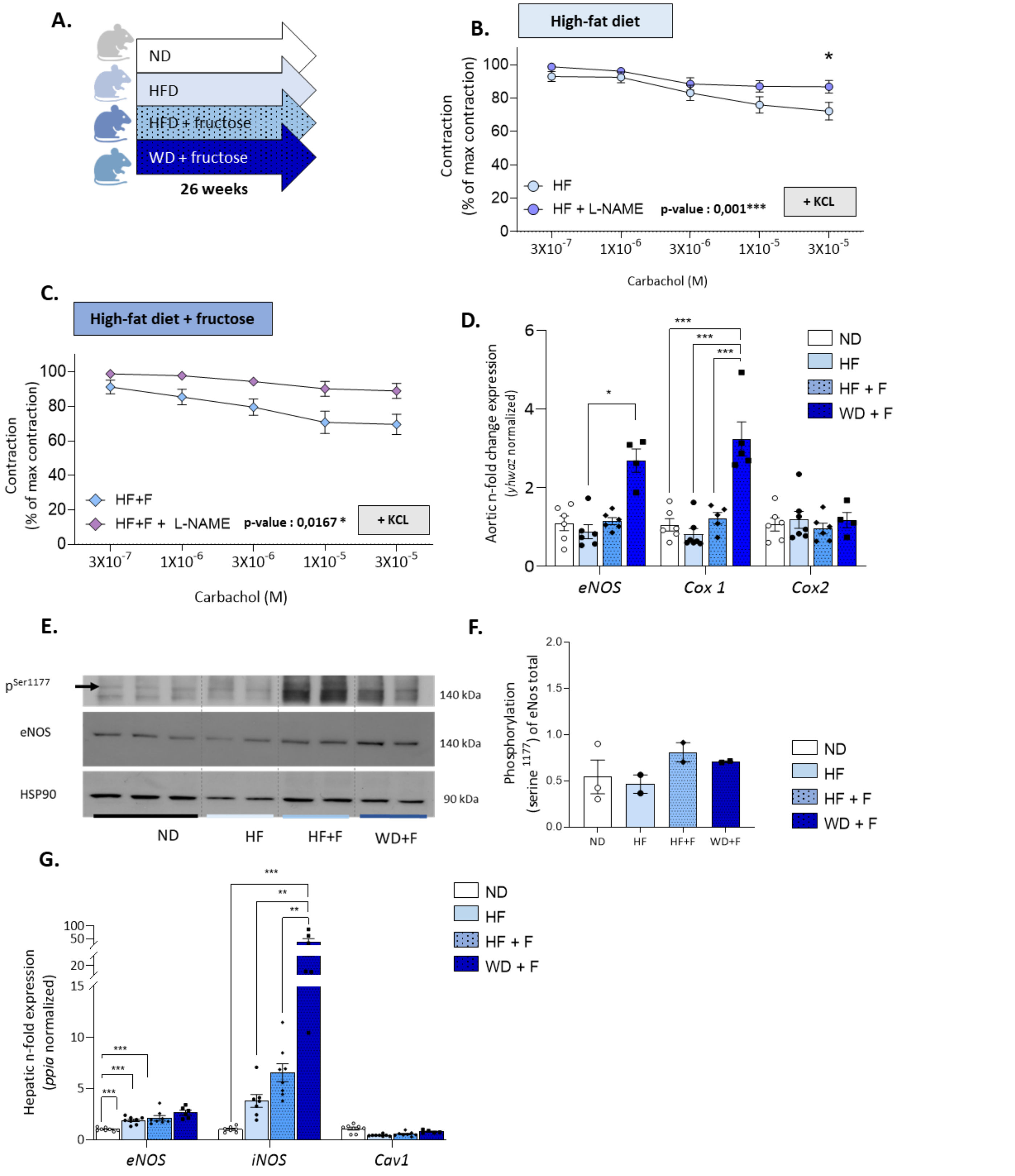


**Supplemental Figure 3**. A. Experimental setup B, C Vasorelaxation in superior mesenteric of HF (B), and H+F (C) induced by KCL (EDHF pathway inhibitor) with or without L-NAME (eNOS inhibitor) treatment. D. Aortic gene expression levels normalized with *Ywhaz, n*-fold expression using ND as reference E. Detection of phosphorylated Serine$^{1177}$ form of eNOS, total eNOS and HSP90 by western blot in aorta of ND, HF, HF+F and WD+F mice. F. Quantification of phosphorylated Serine$^{1177}$ form of eNOS on total eNOS. HSP90 was used as a loading control. G. Hepatic gene expression levels normalized with *Ppia, n*-fold expression using ND as reference. n = 4-8 mice per group; n= 9-15 rings per group ; statistics : Kruskal–Wallis with Dunn's multiple comparisons test (D,F,G); two-way ANOVA followed by post hoc multiple comparisons with Bonferroni correction (B,C). The reported p-value corresponds to the effect of the treatment factor (L-NAME) as determined by two-way ANOVA (B,C). Data are shown as mean +/- SEM, *p<0,05, **p<0,01, ***p<0,001, ****p<0,001. Cav1 : caveolin1, COX 1/2 : cyclooxygenase 1/2; eNOS : endothelial nitric oxide synthase, F : fructose, HbNO : nitrosylated hemoglobin, HF : high-fat, iNOS: inducible nitric oxide synthase, L-NAME : L-N$^{G}$-Nitro arginine methyl ester, Ppia : peptidylprolyl isomerase A, WD : western diet, WT : wild type, ywhaz : 14-3-3-zeta.

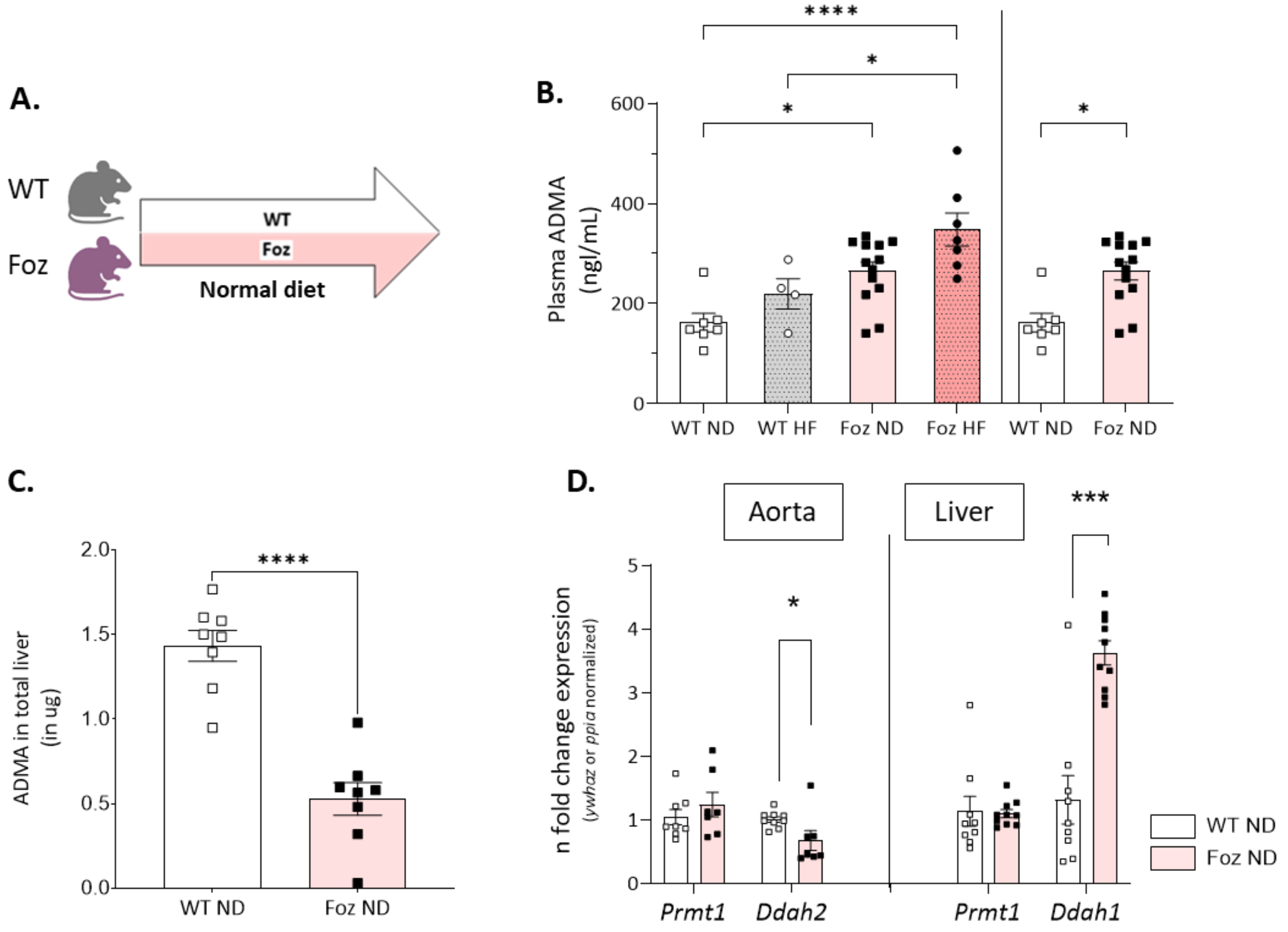


**Supplemental Figure 4**. A. Experimental setup. B. Plasma ADMA levels in ng/ml. C. ADMA content in total liver. D. Aortic and hepatic Prmt1 and Ddha1/2 gene expression levels normalized with Ywhaz and Ppia, respectively, n-fold mRNA expression calculated using WT ND as reference. n=7-11 mice per group ; statistics : unpaired two-tailed t-test (B,C,D), unpaired two-tailed Mann-Whitney test (D). Data are shown as mean +/- SEM, *p<0,05, **p<0,01, ***p<0,001, ****p<0,001. ADMA : asymmetric dimethylarginine, Ddah1/2 : dimethylarginine dimethylaminohydrolase, Ppia : peptidylprolyl isomerase A, Prmt1: protein arginine methyltransferase 1, WT : wild type, ywhaz : 14-3-3-zeta.

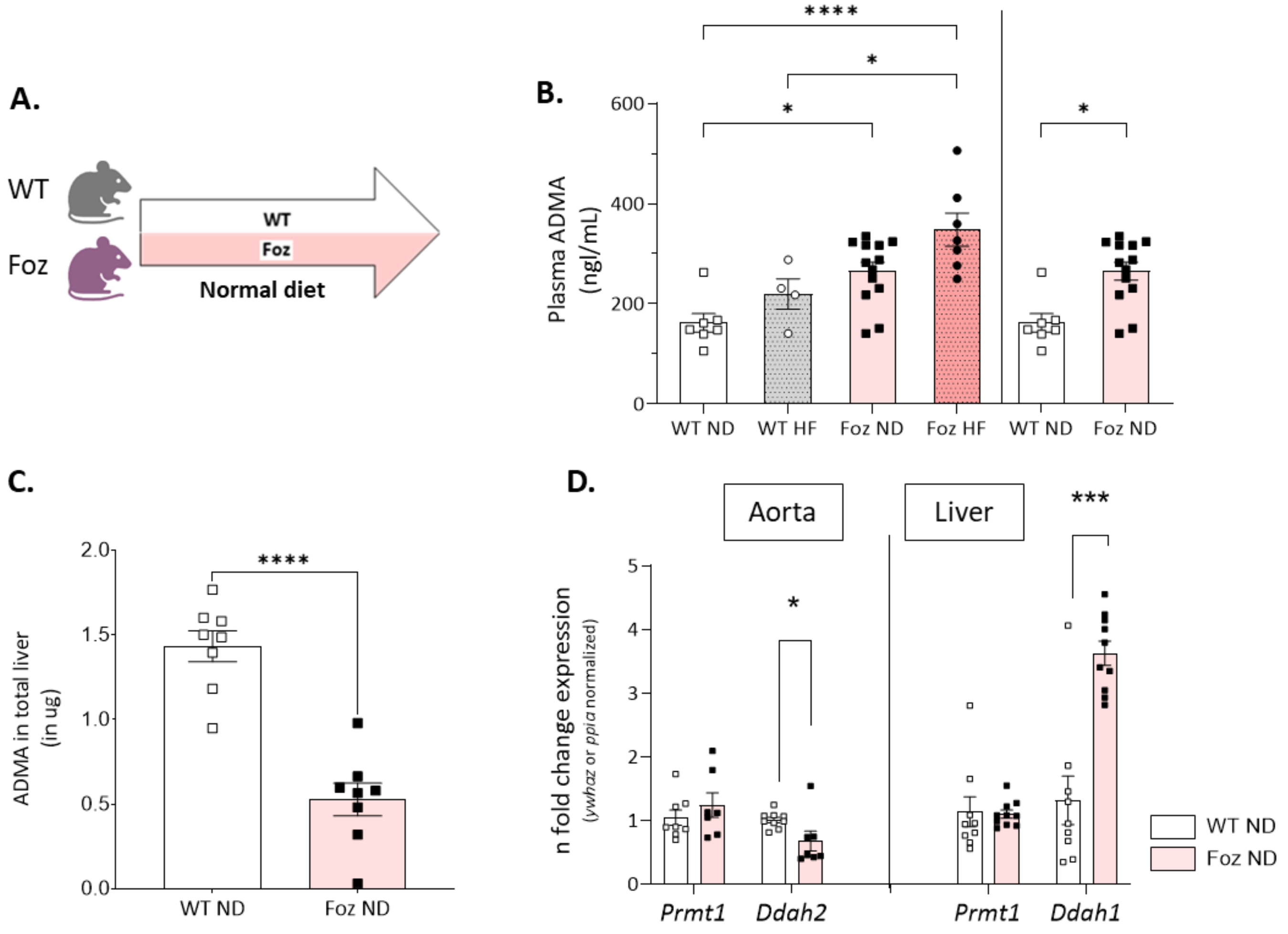


**Supplemental Figure 5.** A. Experimental setup. B. Plasma ADMA levels in ng/ml. C. ADMA content in total liver. D. Aortic and hepatic Prmt1 and Ddha1/2 gene expression levels normalized with Ywhaz and Ppia, respectively, n-fold mRNA expression calculated using WT ND as reference. n=7-11 mice per group ; statistics : unpaired two-tailed t-test (B,C,D), unpaired two-tailed Mann-Whitney test (D). Data are shown as mean +/- SEM, *p<0,05, **p<0,01, ***p<0,001, ****p<0,001. ADMA: asymmetric dimethylarginine, Ddah1/2: dimethylarginine dimethylaminohydrolase, Ppia : peptidylprolyl isomerase A, Prmt1: protein arginine methyltransferase 1, WT : wild type, ywhaz : 14-3-3-zeta.

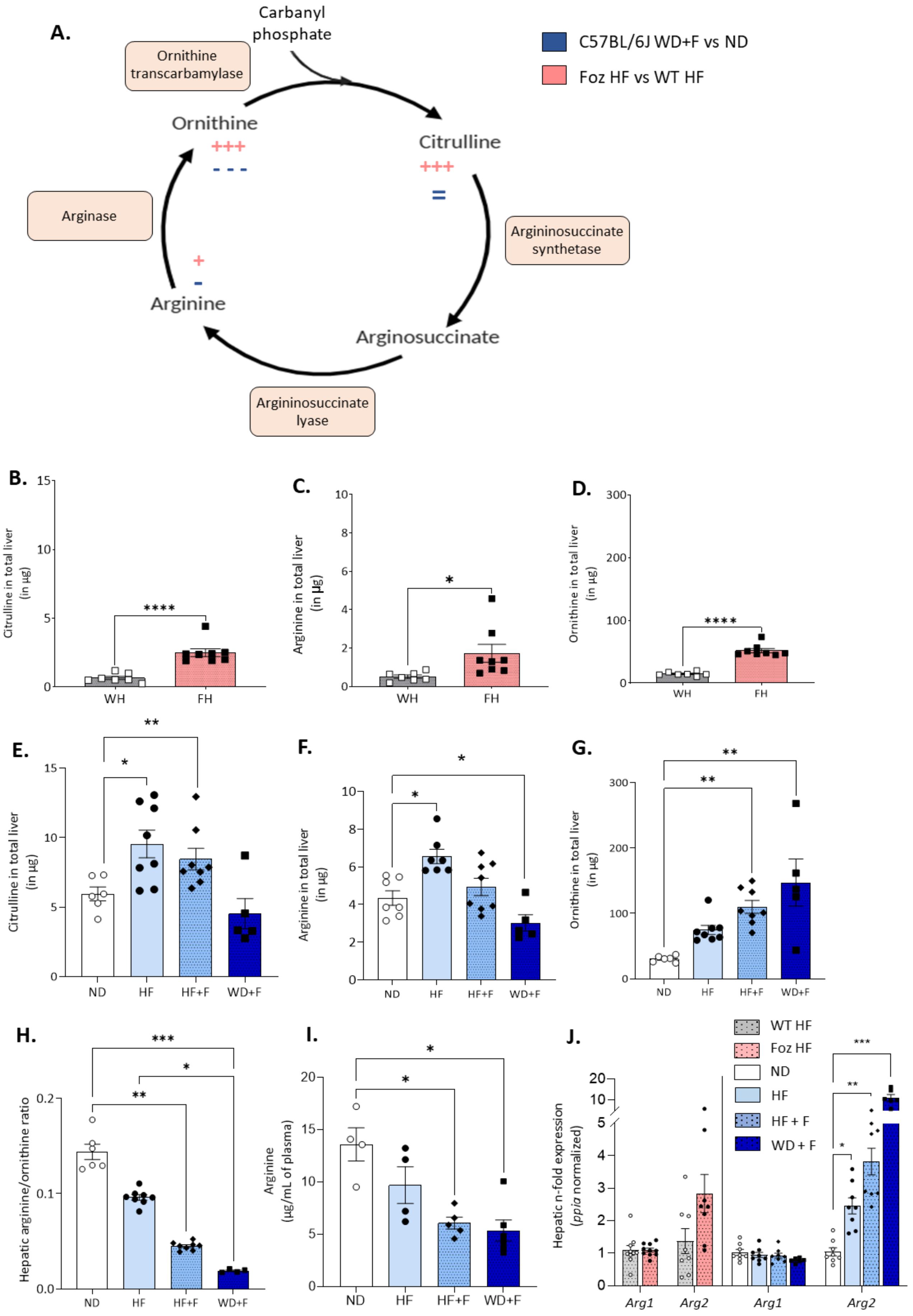


**Supplemental Figure 6.** A.Urea cycle: differences in hepatic metabolite content are depicted in blue for WD + F versus ND mice, and in red for Foz HF versus WT HF mice**. B,C,D.** Citrulline, arginine and ornithine content in total liver in Foz mice compared to WT HF. **E,F,G.** Citrulline, arginine and ornithine content in total liver in ND, HF , HF+F and WD+F mice. **H**. Arginine/ornithine ratio in the liver **I**. Plasma ADMA levels in ng/ml. **J.** Hepatic gene expression levels normalized with *Ppia, n*-fold expression using WT HF or ND as reference. n=4-8 mice per group ; statistics : unpaired two-tailed t-test (B,C,D,J), Kruskal–Wallis with Dunn's multiple comparisons test (E,F,G,I,J) or one-way ANOVA with Tukey's multiple comparisons test (I,J). Data are shown as mean +/- SEM, *p<0,05, **p<0,01, ***p<0,001, ****p<0,001. Arg1/2 : arginase ½ , F: fructose, HF : high-fat, Ppia : peptidylprolyl isomerase A, WD : western diet, WT : wild-type, ywhaz : 14-3-3-zeta.
